# Score matching for differential abundance testing of compositional high-throughput sequencing data

**DOI:** 10.1101/2024.12.05.627006

**Authors:** Johannes Ostner, Hongzhe Li, Christian L. Müller

**Affiliations:** Computational Health Center, Helmholtz Munich, Neuherberg, Germany; Institut für Statistik, Ludwig-Maximilians-Universität München, Munich, Germany; Department of Biostatistics, Epidemiology and Informatics, Perelman School of Medicine, University of Pennsylvania, Philadelphia, PA, USA; Center for Computational Mathematics, Flatiron Institute, New York, NY, USA

**Keywords:** Compositional data, Score matching, Differential abundance, Generative model, Single-cell RNA sequencing, Microbiome

## Abstract

The class of a-b power interaction models, proposed by Yu et al. (2024), provides a general framework for modeling sparse compositional count data with pairwise feature interactions. This class includes many distributions as special cases and enables zero count handling through power transformations, making it especially suitable for modern high-throughput sequencing data with excess zeros, including single-cell RNA-Seq and amplicon sequencing data. Here, we present an extension of this class of models that can include covariate information, allowing for accurate characterization of covariate dependencies in heterogeneous populations. Combining this model with a tailored differential abundance (DA) test leads to a novel DA testing scheme, `cosmoDA`, that can reduce false positive detection caused by correlated features. `cosmoDA` uses the generalized score matching estimation framework for power interaction models Our benchmarks on simulated and real data show that `cosmoDA` can accurately estimate feature interactions in the presence of population heterogeneity and significantly reduces the false discovery rate when testing for differential abundance of correlated features. Finally, `cosmoDA` provides an explicit link to popular Box-Cox-type data transformations and allows to assess the impact of zero replacement and power transformations on downstream differential abundance results. `cosmoDA` is available at https://github.com/bio-datascience/cosmoDA.

## 1 Introduction

Count matrices, detailing the compositional makeup of cellular constituents in a sample, are an important data modality derived from modern high-throughput sequencing (HTS) experiments, including amplicon sequencing (Quinn et al., 2018; Tsilimigras and Fodor, 2016) and single-cell RNA-Sequencing (scRNA-Seq) (Büttner et al., 2021; Heumos et al., 2023). These matrices commonly have the form 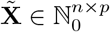 and show the abundance of *p* features (cell types or microbial taxa) in *n* tissues (Regev et al., 2017), bacterial communities (McNulty et al., 2023), or microbial habitats (Turnbaugh et al., 2007). Because sequencing capacity of HTS experiments is technically limited, each sample only represents a small part of a larger population, rendering the sum of counts in a row non-quantitative and making the data compositional (Gloor et al., 2017). Dividing each sample by its total sum yields relative abundance data, which is proportionally equivalent to the original data and constrained to the (*p*−1)-dimensional probability simplex (Aitchison, 1982):

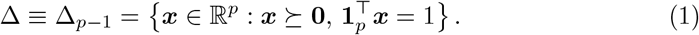

Generative models for HTS-derived compositional data commonly respect compositionality either by transforming the data into Euclidean space through log-ratio or similar transformations (Love et al., 2014; Mishra and Müller, 2022), or by using distributions directly defined on the probability simplex. The Dirichlet distribution is a popular choice due to its relatively simple structure and interpretability (Hijazi and Jernigan, 2009; Wadsworth et al., 2017; Büttner et al., 2021; Ostner et al., 2021). The assumption of independent features (apart from the compositional effect) is, however, a major limitation of the Dirichlet distribution. To allow for more complex dependency structures, Aitchison and Shen (1980) proposed the class of logistic normal distributions, which include the estimation of feature-feature interactions. Several lines of research make successful use of logistic normal models (and extensions thereof) for HTS data (see, e.g., Xia et al. (2013); Zeng et al. (2022)) and address the computational challenges in scaling parameter inference to large-scale datasets (Silverman et al., 2022).

Another challenge in generative HTS data modeling is the presence of zeroes. Since the logistic normal distribution requires the underlying data to be positive due to logarithmic transformations, zero entries in the primary data need to be replaced by positive values (Lubbe et al., 2021; Greenacre et al., 2023). Any such procedure inevitably distorts the measured data compositions, especially for rare features with many zero entries (Lubbe et al., 2021), resulting in another source of modeling inaccuracy.

In his seminal work, Aitchison (1985) provided a general class of distribution, the *A*^*p*−1^ class, that includes the logistic normal and the Dirichlet distribution as special case. This class forms the basis for more recent models that extend the *A*^*p*−1^ class and do not require zero imputation. For low-dimensional data, Scealy and Wood (2022) and Scealy et al. (2024) introduced the polynomially tilted pairwise interaction (PPI) model, which has properties similar to the Dirichlet distribution at the boundaries of the simplex. The class of a-b power interaction models, introduced by Yu et al. (2024), achieves validity on the simplex boundaries by replacing the logarithm with power transformations. These works further use score matching estimation (Hyvärinen, 2005) for computationally efficient parameter inference, reducing the estimation problem to solving a (regularized) quadratic optimization problem. However, both the PPI and the a-b power interaction model currently only allow to model a homogeneous sample population and cannot describe differences between groups of samples.

A central task in HTS data analysis is the detection of significant differences in the feature composition, given environmental, clinical, or host-specific perturbations or variations. This problem, also known as differential abundance (DA) testing, faces the same challenges as generative modeling (Gloor et al., 2017; Tsilimigras and Fodor, 2016). While compositionality and zero handling are discussed in most state-of-the-art DA testing methods (Lin and Peddada, 2020; Zhou et al., 2022; Nearing et al., 2022), only few methods explicitly include interactions between compositional features in their testing procedure (Ma et al., 2024). Such interactions, however, may contribute to the false discovery of certain features that are not directly impacted by the perturbations or covariate changes, but simply strongly correlate with the differentially abundant feature. Consider a composition of five microbial taxa *a, b, c, d, e*, where *a* and *b* have a symbiotic relationship and their abundances are highly correlated (Figure 1a). A treatment now targets taxon *a* and causes a decline in its population. This will in turn cause the abundance of taxon *b* to also decrease, although it was not directly influenced by the treatment. Classical DA testing methods will not be able to discern between these primary and secondary effects caused by the treatment, detecting both *a* and *b* as differentially abundant.

**Fig. 1.**
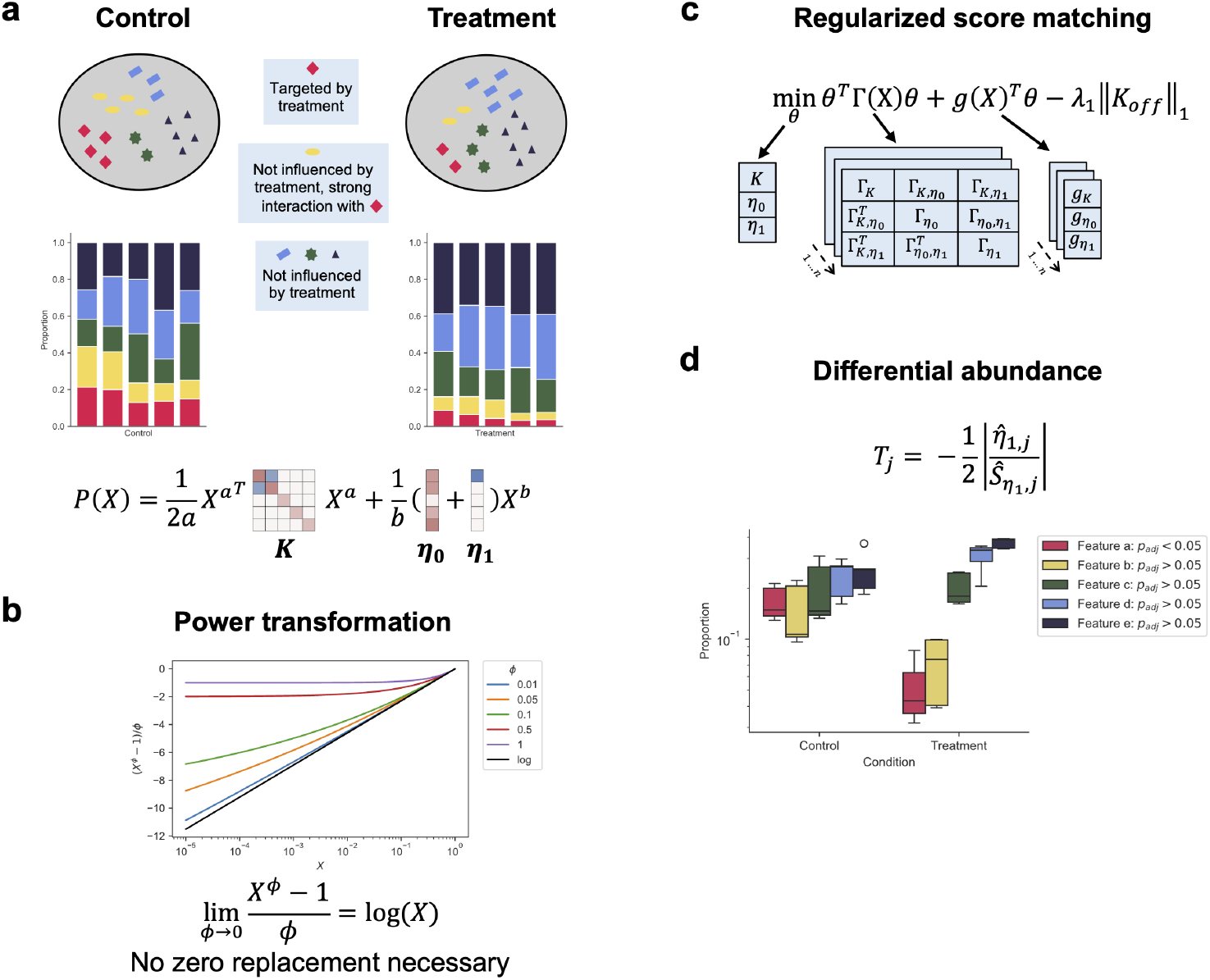
cosmoDAallows to perform generative modeling and differential abundance testing on compositional data with feature interactions. (a) Interactions between features can alter the abundance of features although they are not directly associated with the condition. `cosmoDA`is able to accurately distinguish primary from secondary effects by inferring pairwise feature interactions in addition to the effects associated with the condition. (b) Power transformations allow to analyze compositional data without imputation of zero values. For decreasing exponents, the Box-Cox transformation converges to the logarithm. (c) `cosmoDA`uses regularized score matching for parameter inference. The optimization problem therefore reduces to a quadratic function with parameters Γ and *g* defined by averaging over all samples. (d) Differential abundance testing in `cosmoDA`uses a studentized test statistic. Only the feature primarily associated with the condition (Feature a) has a small adjusted p-value.

In this work, we present a new DA framework, termed `cosmoDA` (**co**mpositional **s**core **m**atching **o**ptimization for **D**ifferential **A**bundance analysis), that addresses the challenge of feature interactions in DA testing. `cosmoDA` is based on the a-b power interaction models from Yu et al. (2024) and introduces a linear covariate effect on the location vector, thus enabling the inclusion of sample group indicators or continuous covariates of interest. We provide a framework for assessing the significance of the estimated covariate effects, which, in the case of group indicator variables, allows principled compositional differential abundance testing. A similar covariate-extended model was introduced for low-dimensional compositional data by Billheimer et al. (2001), albeit only for the special case of the logistic normal model. In the a-b power interaction models, maximum likelihood estimation is not possible due to the intractability of computing the normalzing constant. We thus resort to the score matching framework (Hyvärinen, 2005). By carefully studying the structural properties of the underlying score matching objective, our extended estimation framework retains the favorable quadratic nature of the underlying optimization problem with negligible computational overhead. Regularization on the interaction effects further ensures model identifiability and selection of the most important correlation patterns. The characteristics of the a-b power interaction model thus ensure that feature interactions are adequately considered and zero entries in the underlying data do not need to be replaced or imputed.

The remainder of the paper is structured as follows. In the next section, we introduce `cosmoDA` as an extension of the a-b power interaction model, describe the score matching estimation framework, and explain how the power transformation makes zero replacement obsolete. We then describe the model regularization framework, sketch the computational implementation, and introduce the differential abundance testing framework `cosmoDA`. Section 3 provides several simulated data benchmarks that show-case the ability of `cosmoDA` to (i) correctly estimate sparse interaction matrices in the presence of covariates and (ii) reduce the false discovery rate in differential abundance testing compared to other state-of-the-art methods. We investigate the impact of different power transformations on differential abundance in real scRNA-seq and 16S rRNA sequencing data in section 4 and provide a data-driven method to select the power exponents in practice. Section 5 discusses the results, highlights strengths and limitations of the work, and provides guidelines for future research. `cosmoDA` is available as a Python package at https://github.com/bio-datascience/cosmoDA.

## 2 Methods

We consider to model compositional matrices ***X*** where each row ***x***^(*i*)^ ∈ Δ, *i* = 1, …, *n*, represents a sample, and each column ***x***_*j*_, *j* = 1, …, *p*, represents the *j*th compositional feature. We are motivated by the large number of available biological “compositional” count matrices 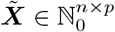 derived from HTS experiments. Important instances include 16S rRNA amplicon sequencing data, where each feature represents read counts associated with a microbial taxon Gloor et al. (2017) and scRNA-Seq experiments, where each feature represents a certain cell-type proportion, as derived from clustered transcriptional profiles (Büttner et al., 2021; Heumos et al., 2023). Due to the compositional nature of the derived count data, a common approach is to scale each observation 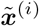 by its library size 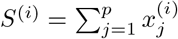 to obtain relative abundancesamples 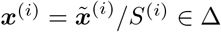 (Gloor et al., 2017).

### 2.1 The covariate-extended a-b power interaction model

Following the proposal in Yu et al. (2024), we model samples in ***X*** through the *a-b power interaction model* on the (*p* − 1)-dimensional simplex Δ. The unnormalized probability density for one sample ***x*** ≡ ***x***^(*i*)^ reads:

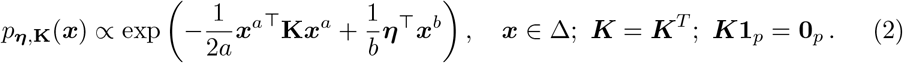

Here, interactions between features are modeled through the *interaction matrix* ***K*** ∈ ℝ^*p*×*p*^, and the *location vector* ***η*** ∈ ℝ^*p*^ describes the base composition of the individual features. This model belongs to the class of exponential family models. Using the conventions in Yu et al. (2024), ***x***^*a*^ ≡ log(***x***); 1*/a* ≡ 1 if *a* = 0, and ***x***^*b*^ ≡ log(***x***); 1*/b*≡ 1 if *b* = 0, power interaction models encapsulate several probability distributions as special cases.

With parameter settings *a* = *b* = 0, the model includes the Dirichlet distribution with the additional constraints ***K*** = 0, *η* ≻ −1, the logistic normal distribution (Aitchison and Shen, 1980) with the constraints 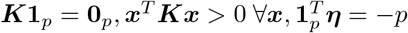,and Aitchison’s *A*^*p*−1^ family of distributions Aitchison (1985) with ***x***^*T*^ ***Kx*** > 0 ***x, η***⪰−1 For the logistic normal case, the interaction matrix ***K*** is equivalent to the inverse covariance matrix of logratio-transformed data, given specific linear transformations (Erb, 2020). With parameter settings *a* = 1 and *b* = 0, the model is equivalent to the PPI distribution (Scealy and Wood, 2022; Scealy et al., 2024) (see Appendix A), and with parameter settings *a* = *b* = 1, the power interaction model is equivalent to the maximum entropy distribution on the simplex with the constraints ***K*1**_*p*_ = **0**_*p*_, ***x***^*T*^ ***Kx*** > 0 ∀***x***, as derived in Weistuch et al. (2022).

As stated in Theorem 1 from Yu et al. (2024), the probability density in Eq. (2) is proper if either

- *a* = 0, *b* > 0,
- *a* = 0, *b* > 0, *η*_*j*_>−1∀_*j*_
- *a* = 0, *b* = 0, log(***x***)^*T*^ ***K*** log(***x***) ≥ 0 ∀***x*** ∈ Δ.
- *a* = 0, *b* > 0, log(***x***)^*T*^ ***K*** log(***x***) ≥ 0 ∀***x*** ∈ Δ.

We next extend the original proposal of the a-b power interaction model by including a (continuous or binary) covariate vector ***y*** ∈ ℝ^*n*^ (or ***y***∈ ***{***0, 1} ^*n*^, respectively) in the model. The covariate describes, e.g., a concurrently measured quantity of interest for each sample, or, more relevant in our context, a condition-specific indicator vector. We model the influence of *y* ≡ *y*^(*i*)^ on **x**^(*i*)^ by introducing a linear model on the location vector ***η***:

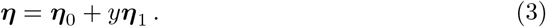

Plugging this model into Eq. (2) results in the covariate-extended a-b power interaction model:

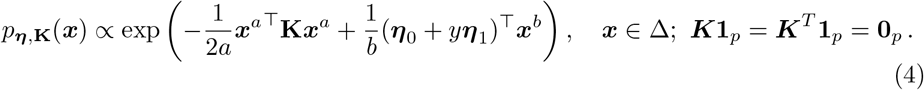

This formulation of the model assumes that all samples stem from an overall population with fixed interaction matrix ***K***, but allows the proportions of features, described by ***η***, to be dependent on the measured covariate *y*. For the probability density of the covariate-extended a-b power interaction model to be proper, the same conditions hold as for the model in Eq. (2), replacing ***η*** with ***η***_0_ + *y****η***_1_.

The model in Eq. (4) forms the basis for our differential abundance testing framework `cosmoDA` (**co**mpositional **s**core **m**atching **o**ptimization for **D**ifferential **A**bundance analysis). In case where ***y*** represents a binary group indicator, e.g., case vs. control samples, `cosmoDA` fits the data to the model and tests for significant changes of the individual components of ***η***_1_. Figure 1 provides a conceptual overview of `cosmoDA`. Before detailing the specific test statistics, we describe the underlying parameter estimation framework.

### 2.2 Model estimation

#### 2.2.1 Score matching for power interaction models

Efficient parameter estimation for the a-b power interaction models (Eq. 2) through generalized score matching (Hyvärinen, 2005, 2007; Yu et al., 2019) was proposed by Yu et al. (2024). Given an (unknown) true data distribution *P*_0_ with density *p*_0_ and a family of distributions of interest 𝒫 (𝒟), score matching tries to find *P* ∈ 𝒫 (𝒟) with density *p* such that the Hyvärinen divergence between the gradients of the logarithm of the densities of *P*_0_ and *P* is minimized:

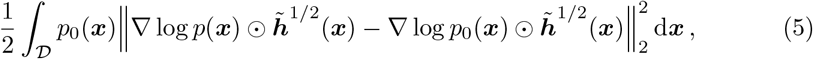

where 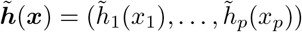 is a weight function. Yu et al. (2022) show that score matching can be performed on domains with positive Lebesgue measure in ℝ^*p*^ by setting 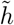 such that 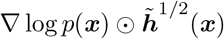 does not vanish at the boundaries of the domain.

Yu et al. (2024) adapted the generalized score matching framework for the a-b power interaction models on the (*p* − 1)-dimensional probability simplex in ℝ^*p*^ by profiling out the last coordinate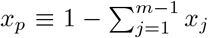 similar to the additive log-ratio transformation. We follow this approach, setting 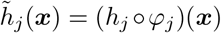 with 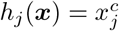 and *φ*_*j*_(***x***) = min{*x*_*j*_, *x*_*p*_, *C*_*j*_} and fixing *C*_*j*_ = 1 and *c* = 2, as recommended by Yu et al. (2024). This results in a weight function 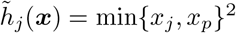. With *p*(***x***) from the family of a-b power interaction models, the following mild assumptions hold (Yu et al., 2024):

1. 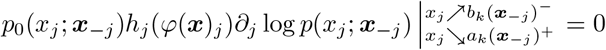 for all *k* = 1, …, *K*_*j*_(***x***_−*j*_) and ***x***_−*j*_ ∈ 𝒮_−*j,𝒟*_ for all *j* = 1, …, *p*;
2. 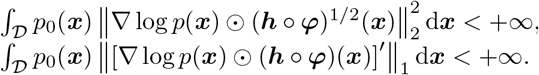
3. ∀*j* = 1, …, *p* and almost everywhere ***x***_−*j*_ ∈ 𝒮_−*j, 𝒟*_,, the component function *h*_*j*_ of ***h*** is absolutely continuous in any bounded sub-interval of the section 𝒞_*j*,𝒟_ (***x***_−*j*_).

Therefore, a consistent estimator of the loss function (Eq. 5) follows as a sample - and feature-wise sum over the entire dataset:

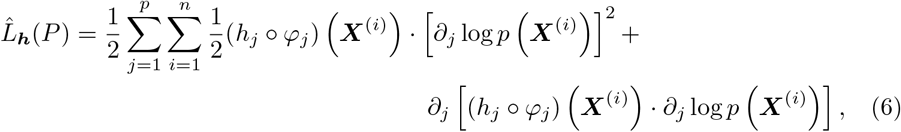

where ***X***^(*i*)^, 1 ≤ *i* ≤ *n*, form an i.i.d. sample from the unknown data distribution *P*_0_ and *P* is an a-b power interaction model with unnormalized density as described in Eq. (2). Aggregating ***K*** and ***η*** to ***θ*** = (vec(***K***), ***η***) and defining *P*_***θ***_ and its density *p*_***θ***_ accordingly shows that the power interaction model without covariate (Eq. 2) follows an exponential-family-type model

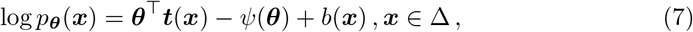

where the function ***t*** (·) denotes the function for the sufficient statistics, *ψ*(·) the cumulant function, and *b*(·) the logarithm of the base measure, respectively.

Then, 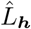 can be reformulated as a quadratic optimization problem:

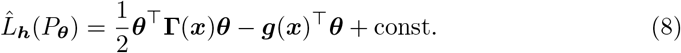

with **Γ**(**x**) ∈ ℝ^*r*×*r*^ and ***g***(***x***) ∈ ℝ^*r*^ are sample averages of known functions in ***x*** only. Analogously, the same considerations hold for the covariate-extended a-b power interaction model (Eq. 4), substituting ***η*** with ***η***_0_ +*y****η***_1_. This substitution does not yet provide individual estimates of ***η***_1_ and ***η***_0_ though, which are required for differential abundance testing. To obtain these individual estimates, a look at the exact derivation of **Γ** and ***g***, as described by Yu et al. (2024), is necessary. We first split the location vector into its two parts ***η***_0_ and ***η***_1_, and set ***θ*** = (vec(***K***), ***η***_**0**_, ***η***_**1**_). After dropping the last coordinate by assuming 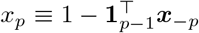 as above, the first and second partial derivatives for the covariate-less model (Eq. 4.1 and 4.2 in Yu et al. (2024)) can easily be adapted to the covariate-extended model:

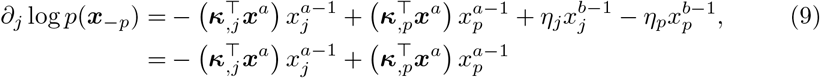

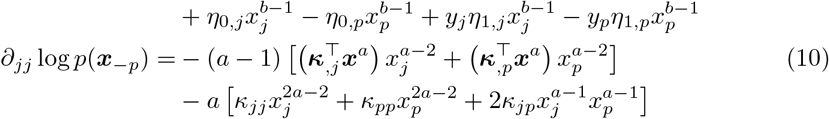

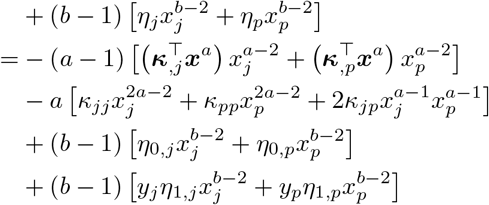

Plugging these definitions into the loss function in Eq. (6) and rearranging the individual terms in the same way as in Yu et al. (2024) yields **Γ** and ***g*** as follows (Figure 1c):

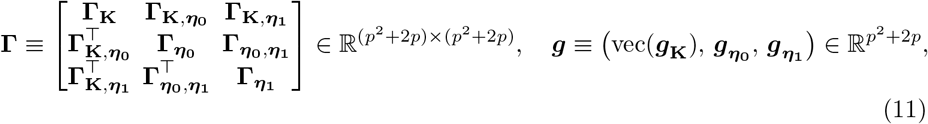

where **Γ** and **g** have a block structure with 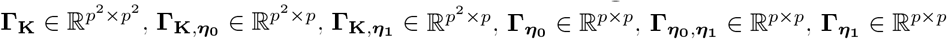, and 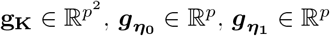.

The exact derivations are shown in Appendix B. By recognizing that each entry of **Γ** and ***g*** can be written as a mean over all samples, the elements related to ***η***_1_ can be computed directly from the elements related to ***η***_0_:

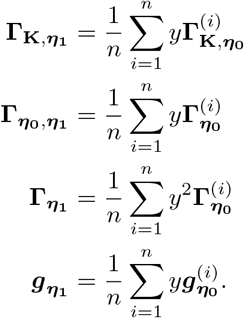

Therefore, the computational overhead for computing the additional sub-matrices and sub-vectors related to ***η***_**1**_ is negligible. Still, the addition of *p* dimensions to the optimization problem (Eq. 8) increases the problem dimensionality from *p*^2^ + *p* to *p*^2^ + 2*p* compared to the covariate-less model.

#### 2.2.2 Model Identifiability through Regularization

Since the number of parameters in the power interaction model scales quadratic with *p*, real HTS data applications are in the high-dimensional regime with more parameters than samples, i.e., *p*^2^ + 2*p* > *n*. To enable model identification, we place a *ℓ*_1_ regularization penalty on the off-diagonal elements **K**_off_ of **K**:

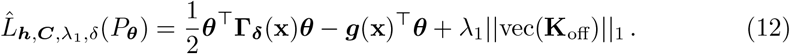

As defined in section 2.2.1, ***θ*** = (vec(***K***), ***η***_**0**_, ***η***_**1**_) comprises all model parameters, and *P*_***θ***_ denotes the power interaction model with probability density *p*_***θ***_ defined in Eq. 4. Following Yu et al. (2024), we multiply the diagonal entries of **Γ**(***x***) corresponding to ***K*** by a factor *δ* > 1 to avoid an unbounded loss function. We denote **Γ**(***x***) scaled diagonal entries as **Γ** _**δ**_ (***x***). Here, we use the default value from the implementation Yu et al. (2024), 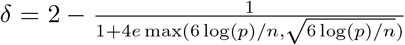. In cases where *p* ≫ *n*, the entries of ***η***_0_ and ***η***_1_ can be penalized as well with a regularization parameter *λ*_2_:

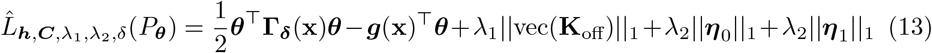

Furthermore, assuming **K** to be sparse matches the widely popular view of sparse association networks between microbial features or cell types (see, e.g., (Kurtz et al., 2015)). In the following, we will focus our attention on models without regularization on the location parameter.

##### Algorithm 1

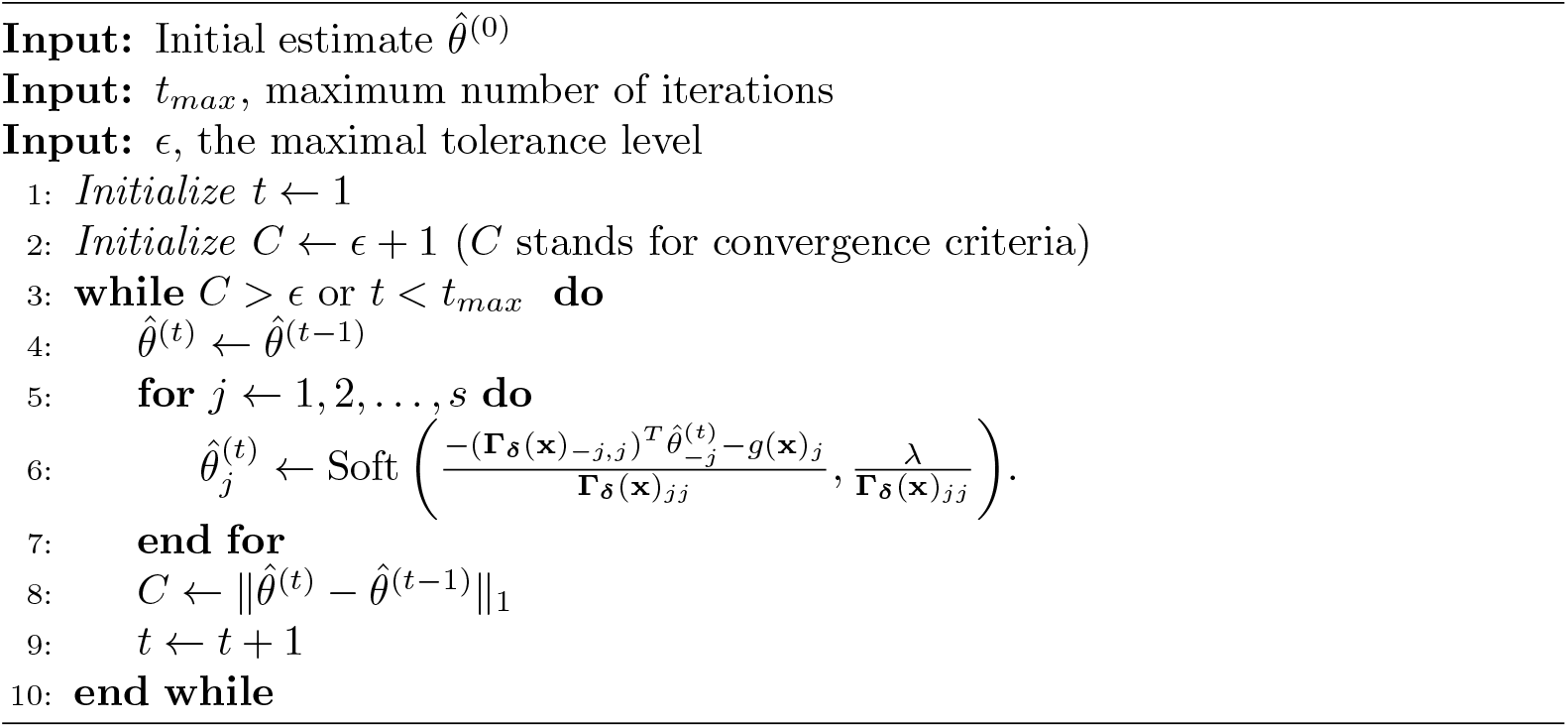

#### 2.2.3 Computational implementation

The regularized score matching loss 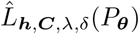 (Eq. 12) represents a (large-scale) *ℓ*_1_-penalized quadratic optimization problem that can be numerically solved with 10 a variety of optimization methods. Here, we follow Yu et al. (2024) and employ a proximal coordinate descent scheme (see also Algorithm 2 in Lin et al. (2016)). This algorithm also covers the covariate-extended a-b power interaction model and is described in Algorithm 1. Here, *s* is the dimensionality of ***θ*** and Soft(·) is the softmax function. The default settings in `cosmoDA` are *ϵ* = 10^−1^ and *t*_*max*_ = 1000.

At its core, our Python implementation of Algorithm 1 uses the C implementation from the *genscore* package Yu et al. (2019, 2024) and included in the `cosmoDA` Python package. The `cosmoDA` package also provides an interface for a-b power interaction models that is equivalent to the R interface in the *genscore* package.

### 2.3 Differential Abundance Testing

One of the key objectives of `cosmoDA` is to determine the statistical significance of the covariate effects ***η***_1,*j*_ for every feature *j* = 1 … *p*. Here, we combine results from Zhou et al. (2022) and Scealy and Wood (2022) to test the null hypothesis

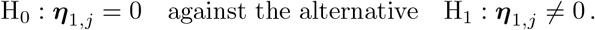

Let 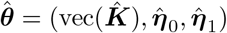 be the parameter estimates obtained from the score matching estimation framework. Scealy and Wood (2022) show that, under certain technical conditions and assumptions (see Theorem 1 and 2 in (Scealy and Wood, 2022)) the quantity 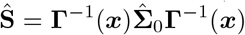 yields a consistent estimator for 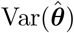.In `cosmoDA`, we estimate 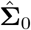 as follows:

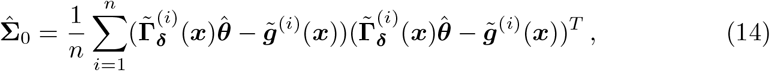

where 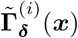 and 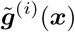 are the components of 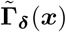 and ***g***(***x***) corresponding to the *i*-th sample. By selecting the components of Ŝ corresponding to *η*_1,*j*_, we derive the studentized test statistic

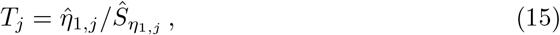

which approximately follows a *t* -distribution with *n*− 3 degrees of freedom. The corresponding asymptotic p-values read:

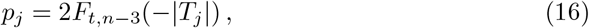

where *F*_*t,n*−3_ is the cumulative distribution function of the *t* -distribution with *n*^−^ 3 degrees of freedom (Zhou et al., 2022). In `cosmoDA`, raw p-values are adjusted for multiple testing using the Benjamini-Hochberg procedure (Benjamini and Hochberg, 1995) (see Figure 1d for illustration).

### 2.4 Model selection and hyperparameter tuning

To make `cosmoDA` fully data-adaptive, we provide several strategies to select the hyper-parameters of the framework. We first describe regularization parameter selection, followed by a novel data-driven approach to select the exponents *a* and *b* in a-b power interaction models.

#### 2.4.2 Regularization parameter selection

`cosmoDA` provides several model selection methods to determine the regularization parameter *λ*_1_ (and *λ*_2_, respectively). The default strategy is k-fold cross-validation (*k* = 5) with the 1SE rule (Hastie et al., 2009). Here, the largest *λ*_1_ value is chosen that lies within one standard error band of the *λ*_1_ that minimizes the cross-validated regularized score matching loss (Eq. 12). The range of the *λ*_1_-path is chosen to cover the whole range of possible sparsity of ***K***, i.e., from a fully dense ***K*** to a diagonal ***K*** (see Fig. E16b). This range depends on the dimensions of the dataset at hand and the chosen power transformation (see Appendix C). Per default, our implementation considers 100 *λ*_1_-values log-linearly spaced in the interval [10^−6^, 1].

`cosmoDA` also allows *λ*_1_ to be selected via the extended Bayesian Information Criterion (eBIC, Foygel and Drton (2010)). Following Yu et al. (2019), the eBIC for the a-b power interaction model reads:

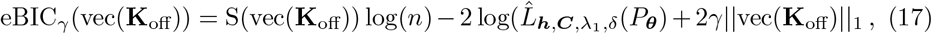

where S(vec(**K**_off_)) denotes the size of the support of vec(**K**_off_). The default *γ* value is *γ* = 0.5.

#### 2.4.2 Data-driven selection of a-b powers

A key strength of a-b power interaction models is their seamless applicability to compositional data with excess zeros. While the limiting case *a* = *b* = 0 (i.e., log-transforming the data) requires a strategy for zero replacement or zero imputation (Lubbe et al., 2021; Greenacre et al., 2023) with potentially detrimental effects for downstream analysis (Te Beest et al., 2021), we propose a data-driven tuning strategy for power interaction models with powers *a* > 0 and *b* > 0 that keeps the original data unaltered. For simplicity, we consider the setting *a* = *b*.

We first note that the power transformations in the models 2 and 4 are similar to the Box-Cox transformation (Box and Cox, 1964) of the form:

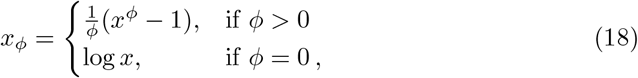

with 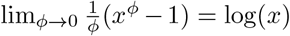(see also Fig. 1b for illustration). The Box-Cox transformation and the power transformation used in a-b power interaction models (Eqs. 2, 4) are, however, not equivalent due to the −1 term in the Box-Cox transformation. By introducing scaling factors for the score matching elements in Eq. 11, we can nevertheless achieve the same asymptotic approximation to the logarithm as the Box-Cox transformation (see Appendix C for details).

While *ϕ* is typically tuned to make the transformed data approach normality, we follow a geometric strategy inspired by the one presented in Greenacre (2024). Specifically, we determine *ϕ* = *a* = *b* to let the resulting “geometry” of transformed data be as similar as possible to the appropriate log-ratio geometry. This is achieved by maximally aligning the principal component (PC) embedding of log-ratio transformed data with imputed zeros and the PC embedding of a power transformation with parameter *ϕ* of the data with zero entries Tsagris et al. (2016). Maximal alignment is defined as the highest Procrustes correlation of both embeddings over a range of values ∈*ϕ*]0, 1[. In the case of power interaction models, it is natural to select a power that closely matches the geometry of data after the additive log-ratio (ALR) transform, since a-b power interaction models with *ϕ* = *a* = *b* = 0 are a generalization of Aitchison’s *A*^*p*−1^ distributions after ALR transformation of the data (Aitchison and Shen, 1980). Since equal dimensionality of the ALR and power-transformed data is required, we append the column 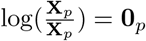 to the ALR transformation of **X** before performing PC analysis.

The original procedure to obtain maximal Procrustes correlation is outlined in Greenacre (2024). We use the same procedure, but with different input matrices. Let ***X***_*ϕ*_ be the Box-Cox-like transformed data with

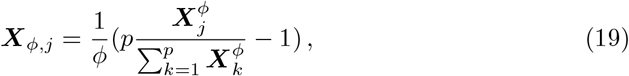

and ***X***_*ALR*_ is the ALR-transformed data (with pseudocount 0.5 for all zeros) with column **0**_*p*_ appended.

We compute the Procrustes correlation *r*_*ϕ*_ between the two data matrices as follows:

(i) Matrix normalization: 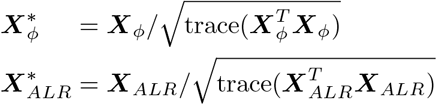
(ii) SVD of cross product: 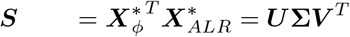
(iii) Optimal rotation matrix: 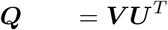
(iv) Sum of squared errors: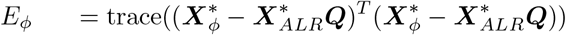
(v) Procrustes correlation: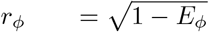

For a given dataset, the optimal power *ϕ*^*^ is determined by *ϕ*^*^ := arg max_*ϕ*_ *r*_*ϕ*_ for *ϕ* ∈]0, 1[.

## 3 Simulation benchmarks

We next provide two simulation studies that benchmark two key features of `cosmoDA`: (i) sparse recovery of feature interactions in the covariate-extended a-b power interaction model and (ii) identification of differentially abundant features in the presence of feature correlations. The first benchmark complements the extensive covariate-free simulation benchmarks of Yu et al. (2024), the second one provides a new realistic semi-synthetic simulation and evaluation setup, incorporating scRNA-Seq data Perez et al. (2022).

### 3.1 Sparse recovery of feature interactions in the presence of a covariate

One of the core strengths of a-b power interaction models is their ability to recover (potentially) sparse feature interaction matrices **K**. Yu et al. (2024) provide an extensive simulation framework that evaluates the influence of hyperparameters, sample size, and interaction topologies on recovery performance of the a-b power interaction model. We focus here on evaluating the influence of covariate inclusion on the model’s ability to identify sparse feature interactions. Specifically, we expect interaction recovery to be *independent of covariate inclusion*.

Following Yu et al. (2024), we generated compositional data 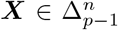 from an model with *p* = 100 features using the model in Eq. 4 with the constraint that ***x***^*T*^ ***Kx*** > 0 ∀***x, η*** ⪰ −1. To probe sample size dependencies, we used two scenarios *n* = 80 and *n* = 1000, respectively. We set ***η***_0_ =− **1**_*p*_, and considered banded interaction matrices **K** with bandwidths *s* = 2 if *n* = 80 and *s* = 7 if *n* = 1000, as suggested by Yu et al. (2024). We further defined the nonzero off-diagonal entries of **K** as **K**_*i,j*_ = |*i*− *j*| */*(*s* + 1) 1 for all *i* = *j*, 1 |*i*− *j*| ≤*s*, and the diagonal entries as the negative sum of the off-diagonals, to ensure the sum-to-zero constraint on the rows of **K** (Figure E2). This definition slightly deviates from the definition in Yu et al. (2024), as the sign of all entries in **K** is flipped, but ensures positive definiteness of **K**. This modification allows the efficient use of the adaptive rejection sampler for data generation, as provided in the *genscore* R package (Yu et al., 2019). For both sample sizes, we generated *R* = 50 replicates of the data.

We applied three different methods for regularized estimation of the underlying interaction matrix **K** to all datasets:

1. The a-b power interaction model (*a* = *b* = 0) without covariate (Eq. 2). This model allows the estimation of **K** and ***η***_**0**_. We estimated these models through the implementation in `cosmoDA`.
2. The covariate-extended a-b power interaction model (*a* = *b* = 0) (Eq. 4). Here, we used a misspecified ***y*** where each entry is drawn uniformly at random from {0, 1}. The model allows the estimation of **K, *η***_0_, and ***η***_1_. We used the implementation in cosmoDA.
3. The graphical lasso model on CLR-transformed data, as introduced in *SPIEC-EASI* (Kurtz et al., 2015). The non-zero entries of the resulting sparse inverse covariance matrix serve as a (mis-specified) proxy for **K**. We used the implementation from the *gglasso* package (Schaipp et al., 2021). Model selection was performed with the extended BIC (eBIC) criterion (Foygel and Drton, 2010) with *γ* = 0.25.

For all three models, we used *n*_*λ*_ = 100 values for the regularization parameter, log-spaced in the range 10^−6^ < *λ*_1_ < 1, and *k* = 5 cross-validation folds. All score matching estimation parameters were set to the defaults recommended by Yu et al. (2024) (see also Section 2.2.3).

To measure recovery performance, we compared the support of the off-diagonal elements of the estimated 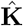 and the ground truth **K** by calculating the true positive rate (TPR) and true negative rate (TNR), and assessing them through Receiver operating characteristic (ROC) curves.

Figure 2 summarizes the average ROC curves for the two different sample sizes. For *n* = 80 (Figure 2a), we observed that both a-b power interaction models showed almost equivalent ability to reconstruct the interaction matrix (mean AUC 0.782 vs. 0.794). Their performance was slightly worse that the graphical lasso (mean AUC 0.806), especially for false positive rates smaller than 0.2. When increasing the sample size to *n* = 1000, all three methods showed improvements in recovery performance, improving the mean average AUC as well as reducing the variance in results (Figure 2b). As expected, including a covariate in the a-b power interaction model had only marginal impact on the mean AUC (0.965 vs. 0.968). Contrary to the low sample size case, both a-b power interaction models significantly outperformed the (misspecified) graphical lasso (mean AUC 0.84) across the entire range of regularization strengths.

**Fig. 2.**
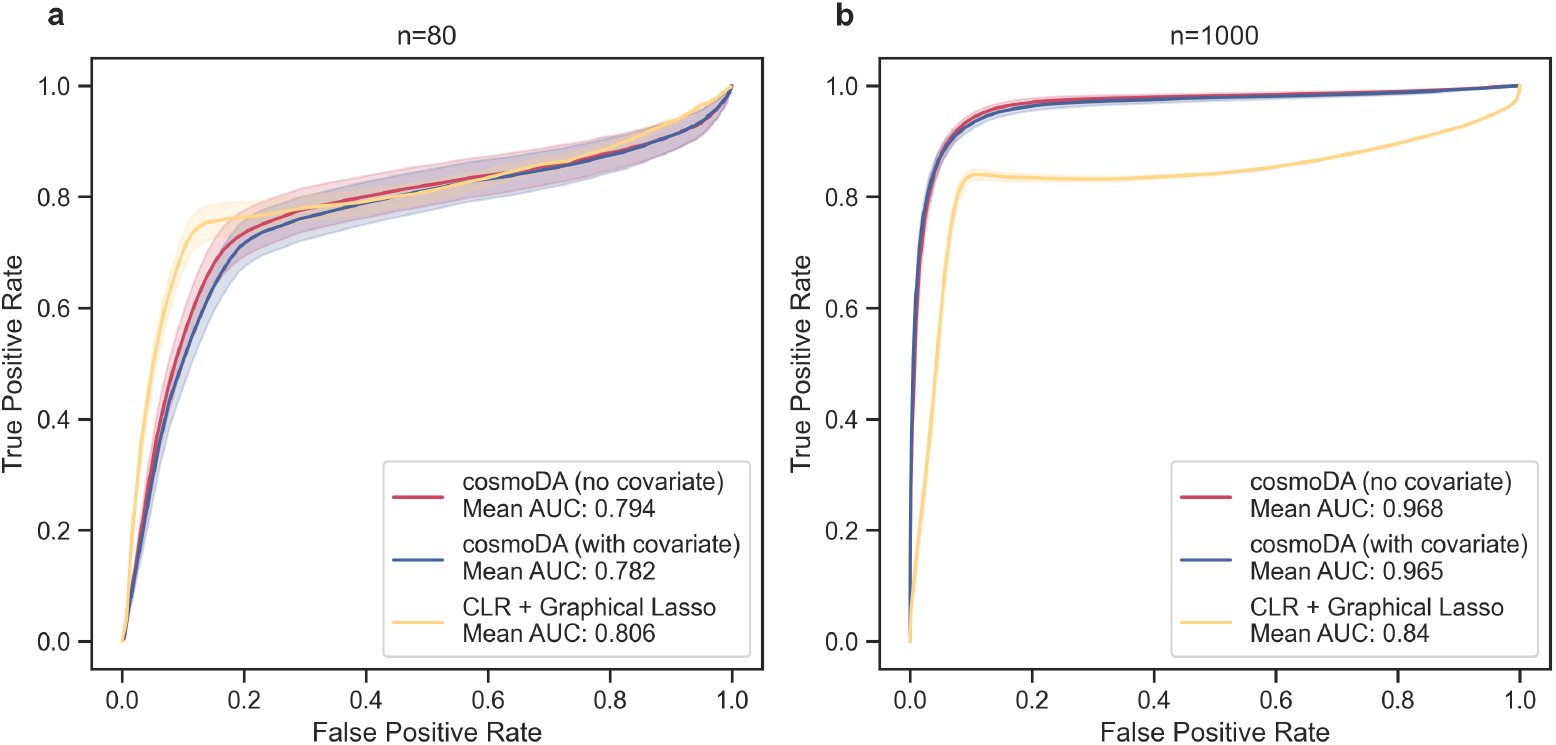
Recovery of K improves with sample size and is not impacted by covariate inclusion. Receiver operating curves for `cosmoDA` with and without covariate effect estimation, as well as CLR transform and graphical lasso for with (a) *n* = 80 and (b) *n* = 1000. The solid lines depict the mean ROC over all 50 generated datasets, the shaded areas show the standard error.

### 3.2 Differential abundance testing in the presence of correlated features

To test the effectiveness of `cosmoDA` in detecting differentially abundant features in the presence of realistic feature interactions, we designed the following semi-synthetic simulation benchmark.

We considered a scRNA-seq data set from Perez et al. (2022) that derived relative abundance values of *p* = 11 types of peripheral blood mononuclear cells (PBMCs) from overall *n* = 352 samples. The samples come from 260 unique subjects, 162 of which are patients with with systemic lupus erythematosus (SLE) (208 samples) and 98 healthy controls (144 samples). We used these data to estimate realistic base values for the interaction matrix **K** and the location vector ***η***, respectively. The base model is the a-b power interaction model without covariate (Eq. 2). We set *a* = *b* = 0 and used *λ*_1_ = 0.043 as sparsity parameter. We considered the NK cell type as the *p*th reference component for all power interaction models due to their high abundance and low variance between groups. The resulting interaction matrix **K**_*B*_ and location vector ***η***_0,*B*_ are shown in Figure 3.

**Fig. 3.**
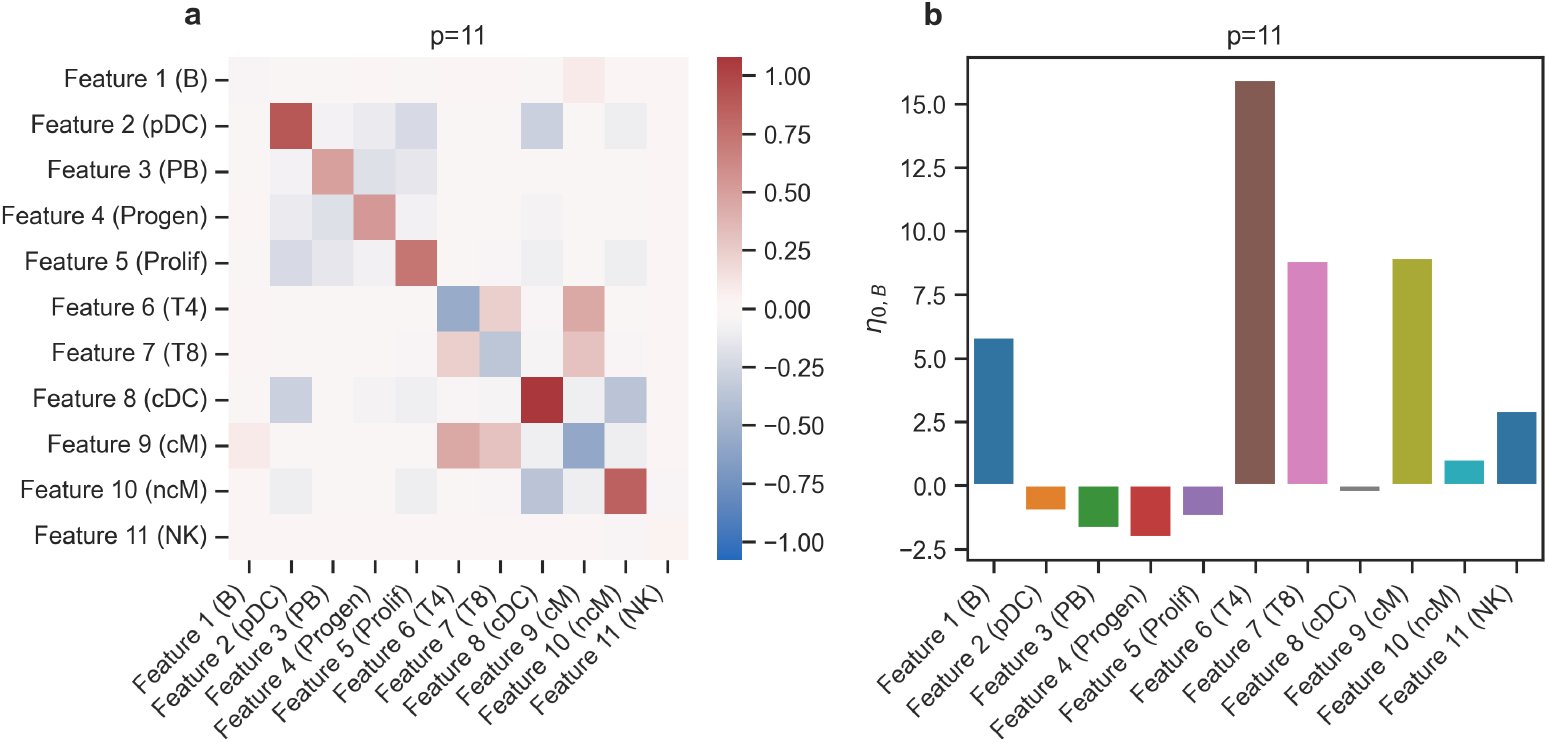
Data generation parameters used for the Differential abundance testing benchmark, *p* = 11.Parameters were generated by running the power interaction model without covariate on the dataset from Perez et al. (2022). The names of the cell types from the original dataset are shown in brackets. (a) Interaction matrix (***K***_*B*_). (b) Location vector (***η***_0,*B*_).

To generate ground-truth differentially abundant cell types, we defined the effect vector ***η***_1,*B*_ = *τ* ***iη***_0,*B*_, where ***i*** is a *p*-dimensional binary vector that indicates the cell types that are influenced by the condition (i.e., are differentially abundant), and *τ* = (− 0.5, −0.3, 0.3, 0.5, 1) controls the relative effect size.

Using this model, we considered three differential abundance scenarios: (i) Estimation when the effect is on a rare cell type with (pDC), (ii) effect estimation on a abundant cell type (T4), and (iii) effects on both cell types (pDC and T4). Two different sample sizes (*n* = 100 and *n* = 1000) were considered for each case. For each of the resulting 30 scenarios, we generated five datasets with *n/*2 control samples (**K** = **K**_*B*_, ***η*** = ***η***_0,*B*_) and *n/*2 case samples (**K** = **K**_*B*_, ***η*** = ***η***_0,*B*_ + ***η***_1,*B*_). To simulate these semi-synthetic data sets, we used the adaptive rejection sampler from the *genscore* R package (Yu et al., 2019).

To showcase the performance of `cosmoDA` for higher dimensional datasets, we conducted another set of simulations with *p* = 99 features. We constructed the corresponding interaction matrix as a block-diagonal matrix, using the original **K**_*B*_ matrix in each of the nine blocks (see Figure E4a). Likewise, we stacked the scenario-specific location vectors ***η***_0,*B*_ and ***η***_1,*B*_ nine times to obtain the high-dimensional location vectors (Figure E4b).

We compared the ability of four different DA testing methods to recover differentially abundant features at an expected FDR level of *α* = 0.05:

- DA testing with `cosmoDA` (*a* = *b* = 0). We used *n*_*λ*_ = 20 values between 10^−3^ and 2 for *λ*_1_ and 5-fold cross validation with the 1SE rule for model selection. All other parameters were set to default values described in Section 2.2.3).
- ANCOM-BC (Lin and Peddada, 2020) with default parameters as an example for a common DA testing method without feature interactions. Since ANCOM-BC assumes count data instead of relative abundances, we scaled the simulated data by the median sequencing depth over all samples in the original dataset and rounded to the nearest integer to obtain comparable counts.
- A Dirichlet regression model and subsequent significance test on the regression coefficients, as implemented by Maier (2014). This model serves as a simple baseline that does not respect feature interactions.
- CompDA (Ma et al., 2024), a recent DA testing method for compositional data, respecting feature interactions via conditional dependency modeling.

Figure 4 summarizes the results for the simulation scenario with the original number of features (*p* = 11). Here, `cosmoDA` showed the overall best ability to recover the true effects (Matthews’ correlation coefficient, Figure 4a), especially when the sample size was larger and for the more abundant cell type T4 (see Figure E5). Importantly, `cosmoDA` showed the lowest FDR value in all scenarios (Figure 4b). Although `cosmoDA` was not able to control the FDR at the expected level of 0.05 in every scenario, the methods without consideration of interactions (ANCOM-BC and Dirichlet) detected more false positive features, with FDR levels averaging between 0.2 and 0.7 in most scenarios. Surprisingly, CompDA did not achieve lower FDR values than ANCOM-BC and performed worse than Dirichlet regression in all cases. We observed slightly elevated FDR levels of `cosmoDA` in cases where the DA cell types were not detected, resulting in FDR values of 1 where one feature was falsely discovered (see Figure E6). While Dirichlet regression and ANCOM-BC struggled with FDR control in all scenarios (see Figure E6), CompDA produced much higher FDR values for the abundant cell type (T4). For smaller sample sizes (*n* = 100) and small effects, `cosmoDA` was not able to consistently detect the differentially abundant features, resulting in lower power for these scenarios. With increasing sample size, the power of `cosmoDA` was on par with the other methods (Figures 4c, E7).

**Fig. 4.**
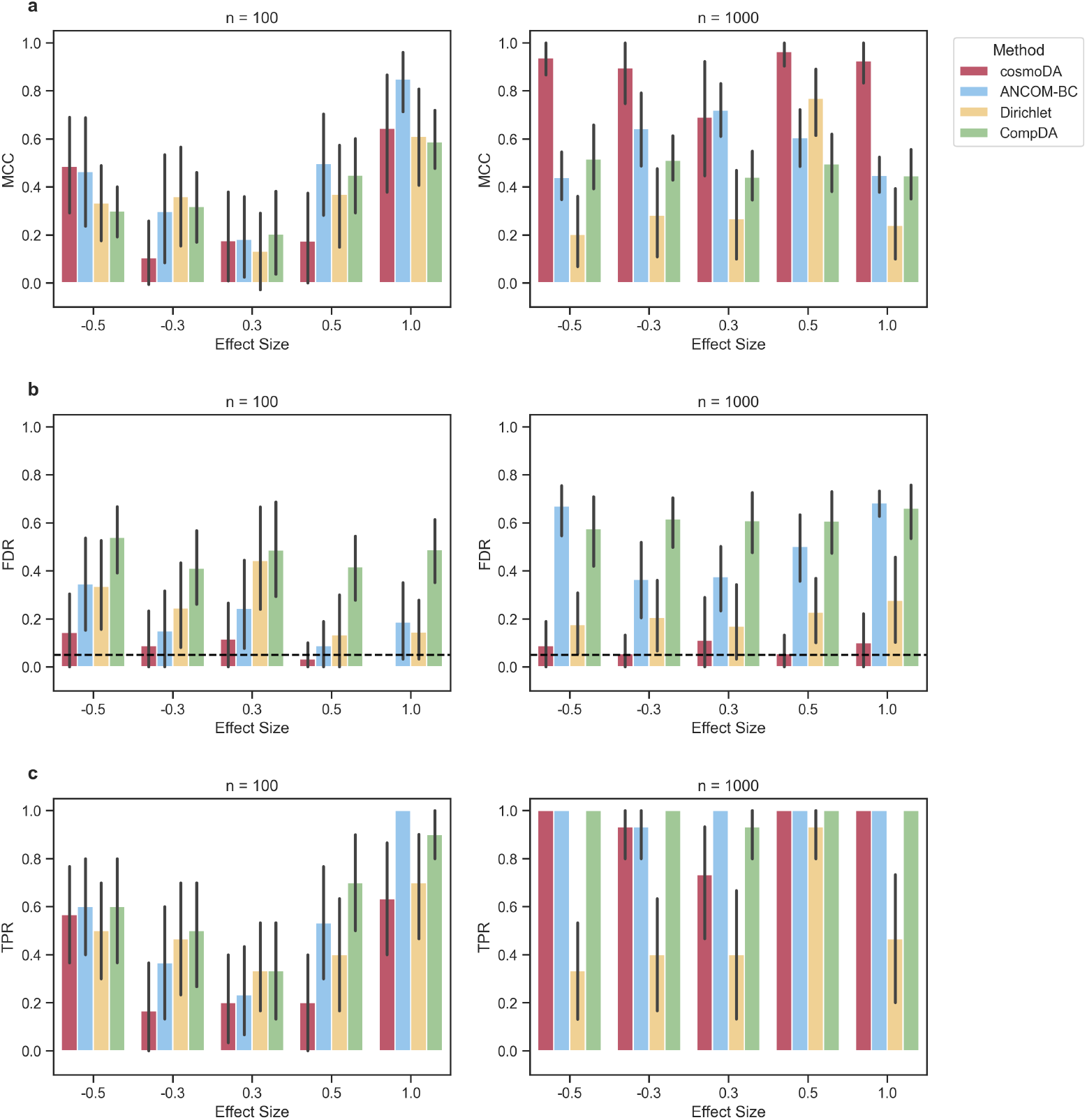
Performance comparison for recovering differentially abundant features across different scenarios, *p* = 11.(a) Matthews’ correlation coefficient. (b) False discovery rate. The dashed line shows the nominal FDR for all methods. (c) True positive rate (power).

In the large-dimensional case (*p* = 99), the performance of all methods decreased in the small sample size scenario, while Matthews’ correlation coefficient was similar to the case of *p* = 11 for larger sample sizes (Figures 5a, E11). Again, `cosmoDA` always showed the lowest FDR, albeit with slightly elevated levels for *n* = 1000, and mean FDR levels between 0.1 and 0.4 for *n* = 100 (Figure 5b). The FDR levels for `cosmoDA` did not show a trend across feature rarity and effect size, while the other methods were not able to produce average FDR levels below 0.5 for effects on rare pDC cells (Figure E12). In terms of power, only ANCOM-BC and Dirichlet regression were able to correctly detect some differentially abundant features for *n* = 100, while all methods showed good power for larger sample sizes (Figure 5c). Breaking these results down by cell type revealed a good power of ANCOM-BC and Dirichlet regression for abundant features, while effects on rare features could not be reliably detected by any method (Figure E13). The unsuitability of `cosmoDA` and CompDA for the case of *p* = 99, *n* = 100 is not surprising, as both models need to estimate pairwise feature interactions in the high-dimensional regime.

**Fig. 5.**
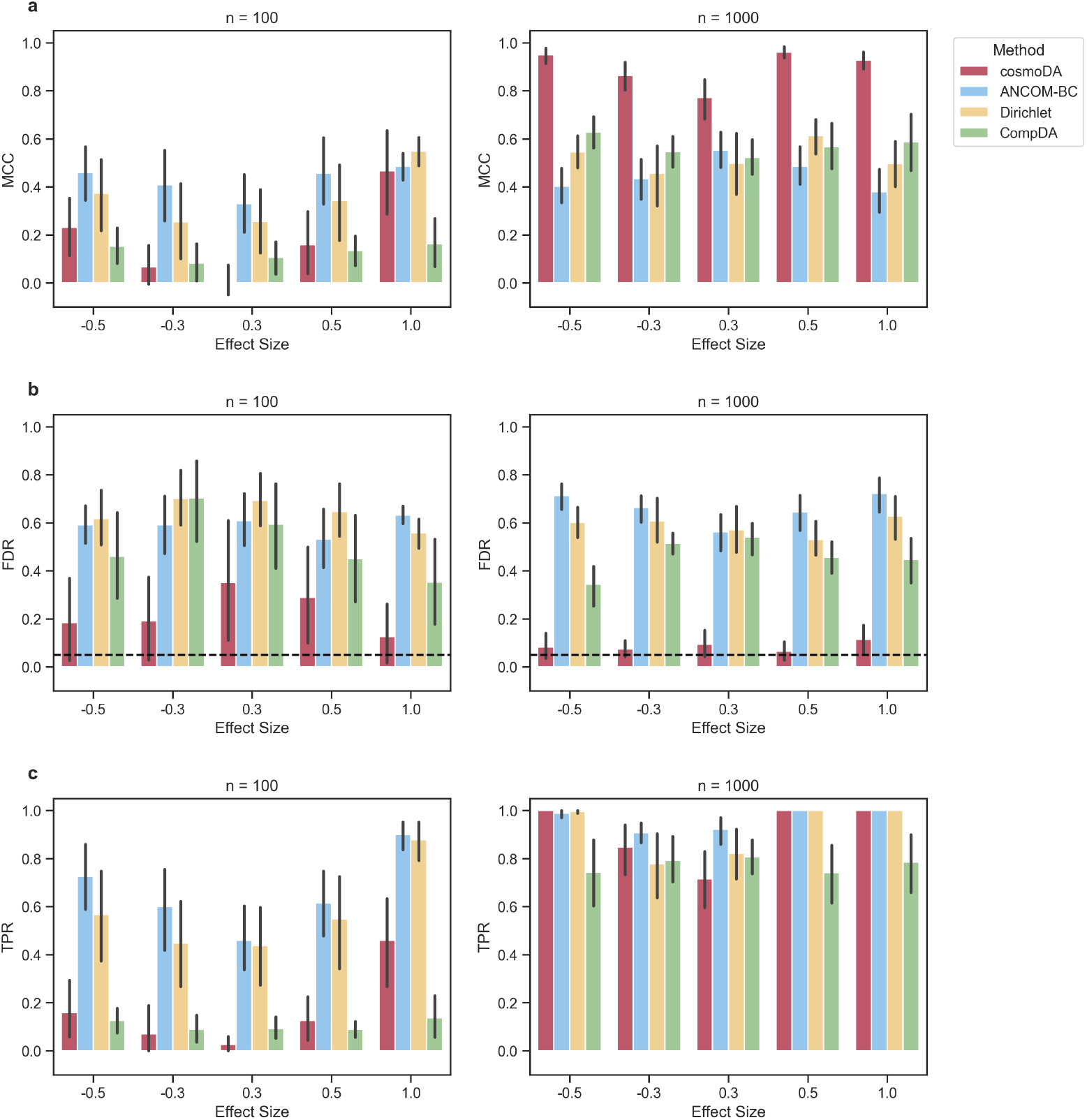
Performance comparison for recovering differentially abundant features across different scenarios, *p* = 99.(a) Matthews’ correlation coefficient. (b) False discovery rate. The dashed line shows the nominal FDR for all methods. (c) True positive rate (power).

## 4 Applications to single-cell and microbiome data

To showcase the DA testing capabilities of `cosmoDA` on real data, we considered two compositional datasets: PBMC abundances derived from scRNA-seq data of SLE patients (as used in the semi-synthetic benchmarks Perez et al. (2022)) and infant gut microbiome data from 16S rRNA sequencing (Yatsunenko et al., 2012). Apart from comparing the empirical results with other state-of-the-art methods, we also evaluated the impact of power transformations (*a* = *b* = *ϕ* ≠ 0) on the downstream DA results.

### 4.1 DA analysis of cell type compositions in patients with systemic lupus erythematosus

We used the scRNA-Seq-derived PBMC data from Perez et al. (2022) (*n* = 352, *p* = 11, see Section 3.2) to estimate differences in cell type composition between subjects with systemic lupus erythematosus (SLE) (n=208) and healthy controls (n=144).

To tune the parameters in `cosmoDA`, we considered the power values *ϕ* = (0.01, 0.02, 0.03, 0.99), as well as the log-log model (*ϕ* = 0) for comparison. We set the range of *λ*_1_ values between 1.5 and 10^−7^ to ensure full coverage of the range of supports of ***K*** for every value of *ϕ*. As before, we used NK cells as the reference cell type for `cosmoDA` and selected the regularization strength via 5-fold cross-validation with the 1SE rule. We used ANCOM-BC, Dirichlet regression, and CompDA for comparison.

We first investigated the influence of our power value tuning schemes for DA analysis. The Procrustes correlation analysis showed that the ALR-transformed PBMC data (with zeros replaced by a pseudocount of 0.5) and the power-transformed data had the highest alignment for a power of *ϕ*^*^ = 0.22 (see Figure 6a). To investigate the impact of zero replacement and the power transform on differential abundance, we also compared the DA testing results of `cosmoDA` for all values of *ϕ* with and without zero entries (Figure 6c, d). Due to the low number of zeroes (4.5%), the impact of zero imputation was negligible for this dataset, making the adjusted p-values with and without zero imputation almost identical. Below a value of 0.8, the exponent of the power transformation only impacted the differential abundance of CD14+ classical monocytes (cM). For higher exponents, almost all cell types showed no differential abundance. Comparing the results of `cosmoDA` with ANCOM-BC, Dirichlet regression, and CompDA (all implemented as described in section 3.2) showed that all methods selected different sets of differentially abundant cell types (Figure 6b). CompDA produced the most conservative results, only finding four DA cell types at an FDR level of 0.05. On the other hand, Dirichlet regression found all cell types to be differentially abundant. Interestingly, `cosmoDA` was the only method that did *not* select classical monocytes as differentially abundant (Figure 6b). The latter finding is in agreement with a control experiment performed by Perez et al. (2022) that found absolute monocyte abundances to be not differentially abundant in SLE patients.

**Fig. 6.**
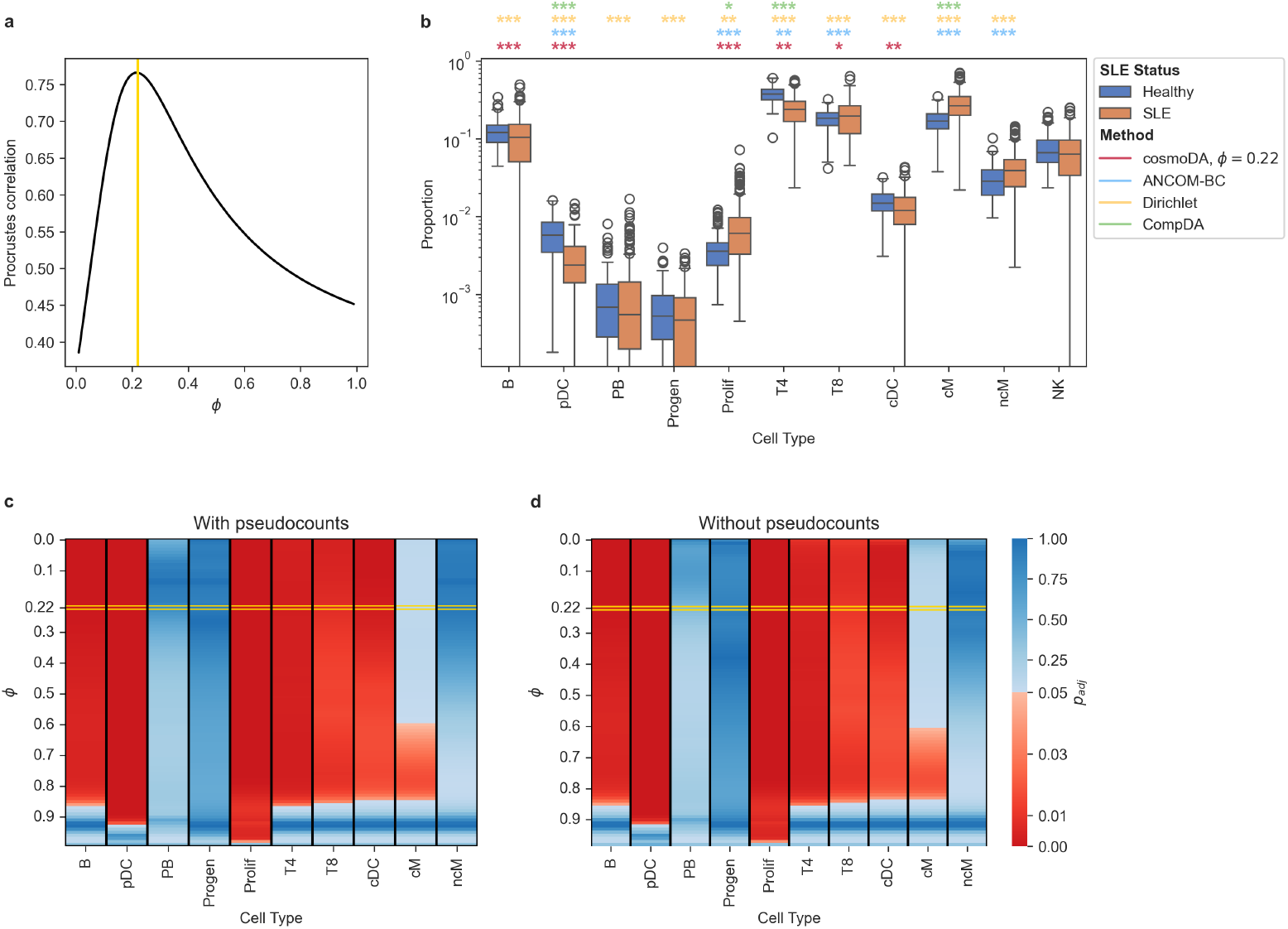
Differential abundance testing with cosmoDA on the lupus dataset. (a) Procrustes correlation between power transformation and ALR transformation with zero replacement. The yellow line (*ϕ*^*^ = 0.22) indicates the maximal Procrustes correlation. (b) Boxplot of relative abundance data without zero replacement. The colored stars indicate the significance level for each method (*: *p*_*adj*_ < 0.05; **: *p*_*adj*_ < 0.01; ***: *p*_*adj*_ < 0.001). Differential abundance results on NK cells (reference in cosmoDA) are omitted. (c) Adjusted p-values for testing differential abundance with `cosmoDA` on zero-replaced data with different power transformations. Red entries denote differential abundance at a level of *α* = 0.05, blue entries denote no differential abundance. The yellow box highlights the adjusted p-values for *ϕ*^*^ determined in a. (d) Same as c, but using the raw data without zero replacement.

### 4.2 DA analysis in microbiome data

To showcase the suitability of `cosmoDA` for microbial 16S rRNA sequencing, we used gut microbiome data from infants in Malawi and the United States (Yatsunenko et al., 2012). We followed the pre-processing in the original ANCOM-BC study Lin and Peddada (2020) and aggregated the data to the Phylum level. We selected all samples from subjects aged less than two years old in Malawi and the United States. Following Lin and Peddada (2020), we next discarded all phyla where more than 90% of samples contained zero entries, resulting in *n* = 97 samples of *p* = 13 phyla. We selected Bacteroidetes as the reference phylum and applied `cosmoDA` with *ϕ* = (0.01, 0.02, 0.03, 0.99), and the log-log model (*ϕ* = 0) to the relative abundances with and without zero replacement. The range of values for *λ*_1_ was set to [10^−12^, 1.5], and we used 5-fold cross-validation with the 1SE rule to select *λ*_1_ for each value of *ϕ*. For this dataset, our power selection scheme identified *ϕ*^*^ = 0.13 to result in the best Procrustes alignment (see Figure 7a). The larger proportion of zero entries in this dataset (28.6%) caused more differences in downstream DA testing results, both on the original and zero-imputed data (Figure 7c, d). While the DA pattern of taxa with no zero entries (Firmicutes, Actinobacteria, Tenericutes, and Proteobacteria) was not impacted by zero imputation, the phyla with at least 20% zero entries (Cyanobacteria, Elusimicrobia, Euryarchaeota, Lentispherae, Spirochaetes, and TM7) were deemed differentially abundant at a smaller power values. Similar to the analysis of the scRNA-seq data, the four DA methods produced different sets of DA taxa at an FDR level of 0.05 (Figure 7b). Dirichlet regression and CompDA seemed to be only sensitive to taxa with high average abundance, while ANCOM-BC and `cosmoDA` were able to also detect differential abundance in rare phyla. The set of DA taxa discovered by `cosmoDA` at *ϕ*^*^ = 0.13 on the data with zero entries was smaller than the set discovered on the same dataset by ANCOM-BC. Nevertheless, `cosmoDA` found multiple phyla that are associated with rural lifestyles (Elusimicrobia, Euryarchaeota, Spirochaetes) to be increased in infants from Malawi (Herlemann et al., 2007; Obregon-Tito et al., 2015). Notably, the ANCOM-BC algorithm involves the replacement of zeros by a small pseudocount (Lin and Peddada, 2020). Indeed, the set of DA phyla discovered by `cosmoDA` on the zero-replaced data with the exponent *ϕ*^*^ = 0.13 (Figure 7c) almost perfectly matched the DA phyla found by ANCOM-BC (except Firmicutes and Proteobacteria). Overall, this confirms that replacement of zero entries in microbial abundance data has significant impact on differential abundance.

**Fig. 7.**
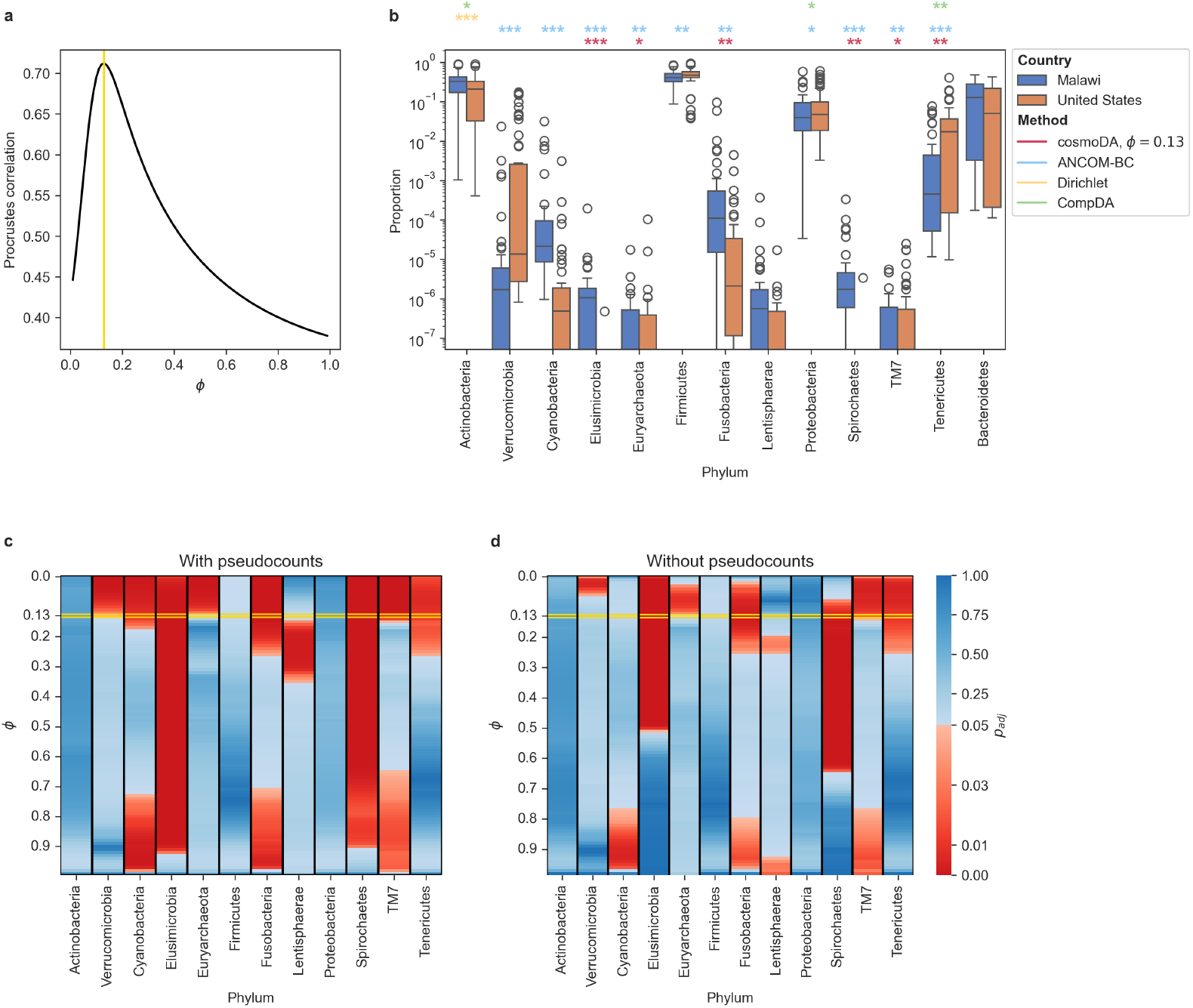
Differential abundance testing (US vs. Malawi) with cosmoDA on infants (age < 2 years) in the the human gut dataset. (a) Procrustes correlation between power transformation and ALR transformation with zero replacement. The yellow line (*ϕ*^*^ = 0.13) indicates the maximal Procrustes correlation. (b) Boxplot of relative abundance data without zero replacement. The colored stars indicate the significance level for each method (*: *p*_*adj*_ < 0.05; **: *p*_*adj*_ < 0.01; ***: *p*_*adj*_ < 0.001). Differential abundance results on Bacteroidetes (reference in cosmoDA) are omitted. (c) Adjusted p-values for testing differential abundance with `cosmoDA` on zero-replaced data with different power transformations. Red entries denote differential abundance at a level of *α* = 0.05, blue entries denote no differential abundance. The yellow box highlights the adjusted p-values for *ϕ*^*^ determined in a. (d) Same as c, but using the raw data without zero replacement.

## 5 Conclusion

Tissues and bacterial communities are complex biological environments, governed by interactions between individual cell types or microbial taxa. The prevailing high-throughput sequencing (HTS) data sets probing these complex mixtures are often compositional in nature. Statistical generative modeling as well as differential abundance testing schemes for such compositional datasets can therefore suffer from inaccuracies if interactions between cells or microbes are not considered in the analysis. Extending the class of a-b power interaction models (Yu et al., 2024) by a linear effect on the location vector, our new method `cosmoDA` allows to accurately model HTS data with pairwise feature interactions in the presence of covariate information. The covariate formulation in `cosmoDA` also seamlessly integrates into the generalized score matching optimization framework (Hyvärinen, 2005; Lin et al., 2016; Yu et al., 2022), facilitating fast and accurate parameter inference. *L*_1_ regularization on the interaction matrix further avoids model complexity explosion and allows to select parsimonious interaction patterns. Compared to the a-b power interaction model without covariates from Yu et al. (2024), the addition of a covariate did not reduce its ability to detect significant feature interactions in our synthetic data simulations. Both the covariate-less and covariate-extended a-b power interaction models outperformed other established procedures for identifying sparse interactions in compositional HTS data when the sample size was sufficiently large.

In the presence of a binary condition, testing for significance of the covariate-related parameters in the location vector acts as a form of differential abundance testing. Here, the parallel estimation of feature interactions helps to avoid false positive detections which are only indirectly related to the condition. In our realistic simulation experiments, `cosmoDA` was the only method to approximately control the false discovery rate in the presence of feature interactions, while no other tested method could distinguish between direct and indirect compositional changes. `cosmoDA` showed reduced power when the sample size was small, but was on par with methods like ANCOM-BC (Lin and Peddada, 2020) for larger numbers of observations. One exception where `cosmoDA` was not able to adequately control the FDR was for misspecified models with more features than samples. We further demonstrated the ability of `cosmoDA` to find biologically meaningful differential abundances on two experimental datasets from human single-cell RNA sequencing and microbiome 16S rRNA sequencing.

The use of power transformations instead of the logarithm in a-b power interaction models allows to keep zero measurements in the data as-is, avoiding distortions caused by imputation of these values. Through a small adjustment in the score matching optimizer, we were able to approximate the log-transformation for exponents approaching zero. Applying `cosmoDA` to real-world single-cell and microbiome datasets, we discovered that zero replacement and the exponent of the power transformation had a considerable impact on downstream DA results in data with excess zeros (Gloor et al., 2017). We further demonstrated that selecting an exponent for the power transformation that approximates the data geometry after an ALR transformation generally produces sensible differential abundance results.

While `cosmoDA` successfully tackles multiple challenges in generative modeling and differential abundance testing, it also has some limitations. Currently, `cosmoDA` can only accommodate a single binary or continuous covariate. Extending the linear model formulation would allow to model more complex scenarios and adjust for confounders in DA testing. For this, the score matching estimator would also have to be extended to multiple covariates. The implementation of such a flexible model could be simplified by using automatic differentiation for determining the elements of **Γ** and ***g*** (Kassel et al., 2024). In addition, we believe that approximation of the logarithm for small exponents can be solved more elegantly by changing the general definition of a-b power interaction models to utilize a true Box-Cox transformation rather than using our proposed adjustments in the score matching optimizer. Estimation of our model also relies on selecting a good reference, which is profiled out in the model formulation. Looping over multiple references and averaging the results, as described by Yu et al. (2024), could avoid this dependency at the cost of computational efficiency.

While we empirically showed the feasibility of `cosmoDA`, we did not provide any guarantees for goodness of fit and convergence. A formal reevaluation and extension of the theoretical considerations provided by Yu et al. (2024) would give more justification to our approach.

Overall, we believe that `cosmoDA` with its abilities to include feature interactions and seamless handling of excess zeros represents a valuable addition to the growing family of differential abundance testing methods. A Python implementation of `cosmoDA` and the power interaction model without covariates is available at https://github.com/bio-datascience/cosmoDA.

## Acknowledgments

We thank Matthias Drton for his inputs to the score matching optimization in `cosmoDA`, and Fabian Schaipp for his recommendations regarding the optimization algorithm. We also thank Oleg Vlasovets for implementing the graphical lasso on the simulated benchmark data. H.L. was supported by NIH grant R01GM123056.

## Author contributions

J.O. developed the idea of `cosmoDA` with help from H.L. and C.L.M. J.O. implemented the method, performed and evaluated all simulations and conducted the data applications. J.O. wrote the manuscript with assistance from C.L.M. All authors read and approved the final manuscript.

## Data and code availability

All datasets used in this article are publicly available. The SLE scRNA-seq data was downloaded from the the Human Cell Atlas platform (GSE174188). The gut microbiome data is stored at MG-RAST https://www.mg-rast.org/index.html under search string “mgp401”, code for data preparation was adapted from https://github.com/FrederickHuangLin/ANCOM-BC. Code for reproducing the analyses in this article is available under https://github.com/bio-datascience/cosmoDA, intermediate data can be found at https://zenodo.org/records/13911623.

## Appendix A Connection between a-b power interaction models and PPI models

The class of polynomially tilted pairwise interaction (PPI) models, introduced by Scealy and Wood (2022), is another class of flexible distributions with feature interactions on the simplex. This class includes distributions of the form

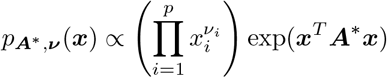

with ***ν*** ≻ −1 ∈ ℝ^*p*^, and ***A***^*^ ∈ ℝ^*p*×*p*^ symmetric with ***A***^*^**1**_***p***_ = 0. Through 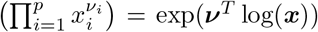, it is easy to see that this class of distributions represents a special case of a-b power interaction models (Eq. 2) with *a* = 1 and *b* = 0Profiling out the last coordinate, i.e 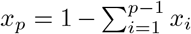, leads to an alternative formulation (Scealy et al., 2024), with parameters ***ν*** ≻ −1 ∈ ℝ^*p*^, ***A***_*L*_ ∈ ℝ^(*p*−1)×(*p*−1)^, and ***c***_*L*_ ∈ ℝ^(*p*−1)^:

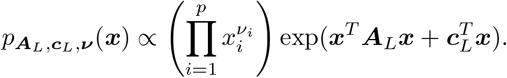

In particular, the transition between the two forms can be achieved by splitting off the last row and column of 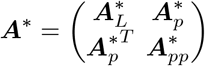. Then 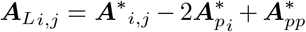 and 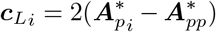. Since ***A***^*^ has one additional parameter, assume 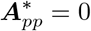 for the reverse transformation. Then, 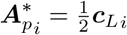 and 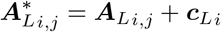.

Applying the equivalent transformations to an a-b power interaction model with *a* = 1 can help with parameter interpretation, as the matrix ***A***_*L*_ usually has full rank.

## Appendix B Derivation of the parameters in the quadratic form of the score matching optimizer

This section details the derivation of the parameters **Γ** and ***g*** in the quadratic formulation of the score matching loss (Eq. 8) and explains their block structure shown in Eq. 11. The elements of ***g*** can be directly derived from the second derivative of log *p*(***x***) (Eq. 10):

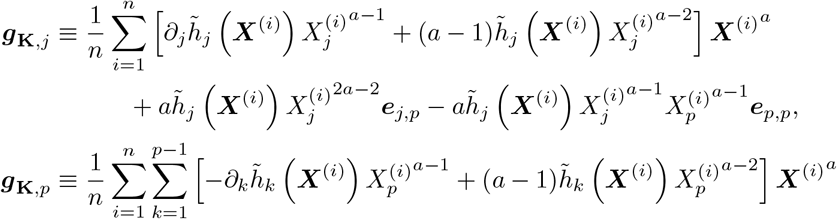

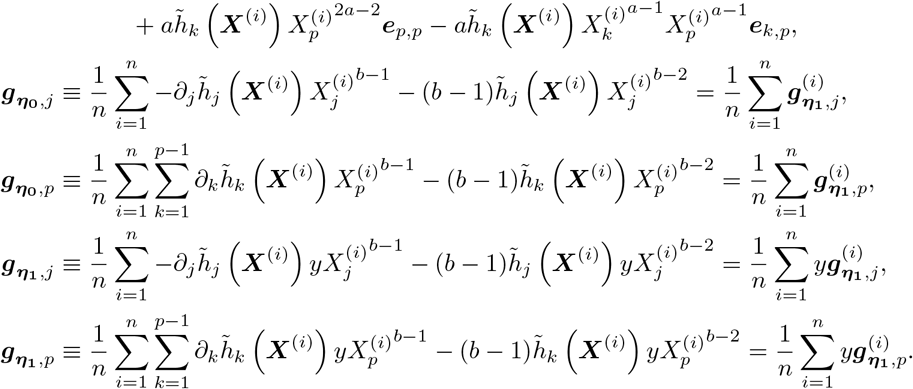

Further, the elements of **Γ** follow from the first derivative of log *p*(***x***) (Eq. 9) and have the same structure as in Yu et al. (2024):

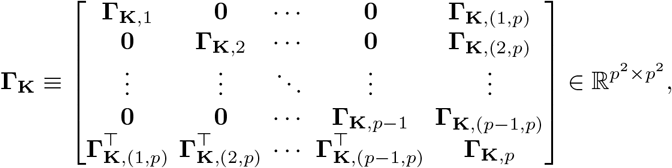

with each block of size p × p, and

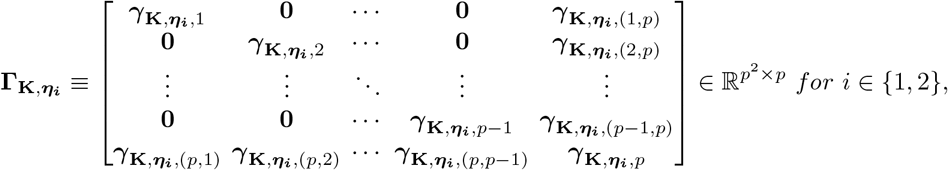

with each block a vector of size p, and

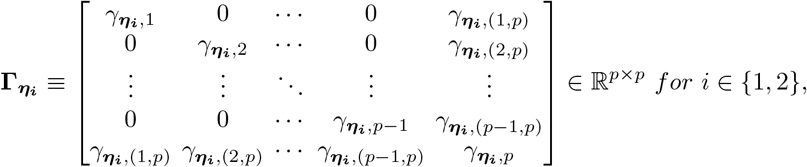

and

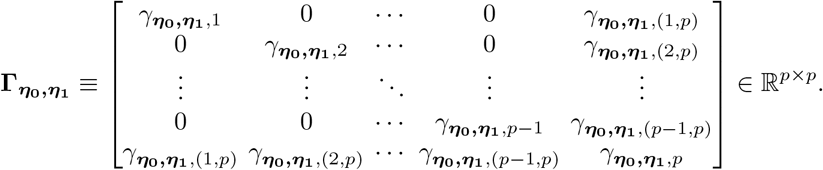

These blocks have the following specific forms. For *j* = 1, …, *p* − 1,

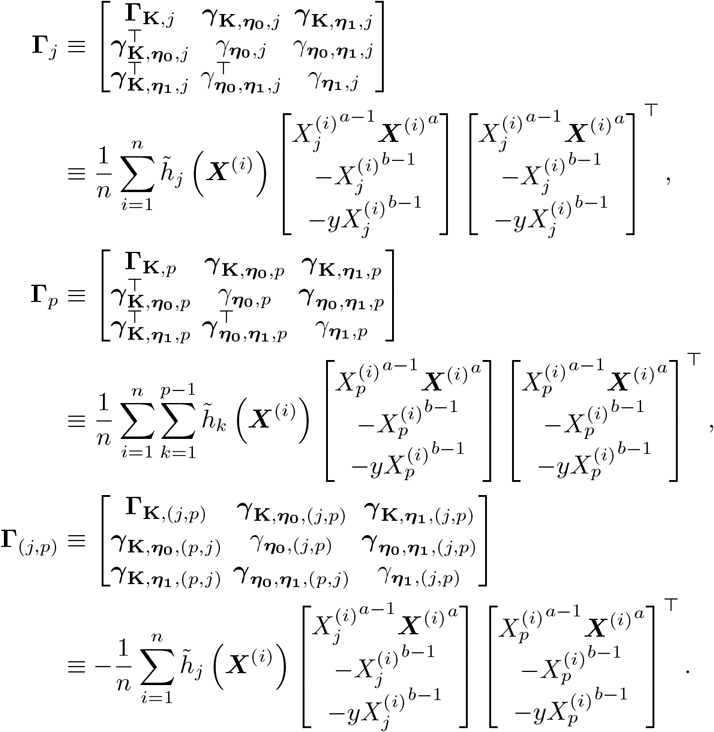

## Appendix C Scaling score matching elements to approximate Box-Cox transformations

As described in Section 2.4.2, the power transformation used for a-b power interaction models (Eqs. 4 and 2) bears striking resemblance to the Box-Cox transformation 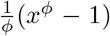. Both transformations are not equivalent though due to the subtraction of 1 in the Box-Cox transformation. This difference causes the a-b power interaction transformation to lose one key property of the Box-Cox transformation - its asymptotic approximation of the logarithm as *ϕ* approaches 0.

Looking at the density of the covariate-extended a-b power interaction model makes this disparity clear:

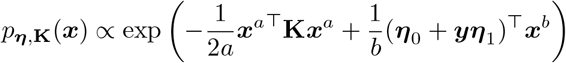

For the terms concerning ***η***_**0**_ and ***η***_**1**_, the subtraction of 1 in the Box-Cox transformation is not dependent on *x* and can therefore be absorbed into the normalizing constant. For the interaction term, replacing ***x***^*a*^ with the Box-Cox transformation in Eqs. 2 or 4 would introduce a scaling factor of order 1*/a*^2^ instead of 1*/a*, leading to a discontinuity of the estimated ***K*** when approaching the log-log model, for which the convention 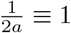 is used (Yu et al., 2024).

We counteract this effect by introducing scaling factors of 1*/a* and 1*/a*^2^ on the components of **Γ** and ***g*** (Eq. 11), based on the matrix multiplication ***θ***^*⊤*^**Γ**(**x**)***θ*** from Eq. 8. In particular, we scale **Γ**_***K***_ by a factor of 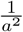 and 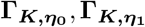, and ***g***_***K***_ by a factor of 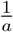 each. This leads to a smooth transition in the estimation of ***K*** when *ϕ* → 0, and also holds for general a-b power interaction models without covariates (Yu et al., 2024).

We showcase the effectiveness of our scaling approach with an example on the scRNA-seq data of SLE patients and healthy controls Perez et al. (2022). For simplicity, we estimate the whole dataset through the covariate-less a-b power interaction model without differentiating between the two groups, use no regularization on the offdiagonal entries of ***K***, and always replace zeros with a value of 0.5. Without the scaling factor, the pattern of the estimated ***K*** approaches the log-solution (*ϕ* = 0), but the scale of the entries is not the same (Figure E14, left column). On the other hand, the entries of ***η*** approach the log-solution also in magnitude (Figure E14, right column). For increasing values of *ϕ*, both the pattern and magnitude of ***K*** and ***η*** gradually diverge, as the power transformation gradually distorts the composition differently.

The median entry of the ratio ***K***_***ϕ*=0**_*/****K***_***ϕ*=*ϕ***_^***′***^ also does not approach 1 as *ϕ* →0 (Figure E15, bottom right). Looking at the components of **Γ** and ***g***, one can see that the median entry of the above ratio follows a log-linear trend for larger values of *ϕ*, but not for smaller exponents if the component is associated with ***K*** (Figure E15, other panels). The scaling factors introduced above correct this trend, such that the ratio is log-linear across the full spectrum of *ϕ*. This causes the estimated ***K*** to approach the solution for *ϕ* = 0 in magnitude (Figure E15, bottom right) without impacting the estimated interaction pattern (Figure E14, middle column) or the estimation of ***η*** (Figure E14, right column).

When combining regularization and power transforms, the dependency between *ϕ* and the scale of entries in ***K*** will lead to different optimal regularization strengths for different exponents (Figure E16a). In fact, a larger exponent and therefore larger scale of ***K*** will require smaller values of *λ*_1_ to cover the whole range between ***K*** with full support and a diagonal ***K*** (Figure E16b). Therefore, the range of values for *λ*_1_ should always be adapted to the current data and power transform.

## Appendix D Testing for differential abundance without feature interactions

We also compared the methods on simulated data without feature interactions to show the suitability of `cosmoDA` if no significant feature associations are present. To this end, we applied the a-b power model solution with *a* = *b* = 0 and *λ*_1_ = 2 to the dataset from Perez et al. (2022), resulting in ground truth parameters of *K*_*B*_ = **0**^11×11^, and ***η***_0,*B*_ as shown in Figure E13. We used the same setup as before to select differentially abundant cell types and effect sizes and again chose *n* = 100 and *n* = 1000, simulating five replicates for each of the 30 scenarios as described above.

If no significant feature interactions were simulated, `cosmoDA` and CompDA showed similar overall performance as before, while the MCC of ANCOM-BC and Dirichlet regression significantly improved (Figures D1a, E8). This improvement was due to a reduction in falsely discovered effects by these methods (Figure D1a), which shows that the high FDR of ANCOM-BC and Dirichlet regression in the previous simulation were caused by secondary effects due to feature interactions. In terms of power, all methods showed similar strength as before (Figures D1c, E10). Nevertheless, `cosmoDA` was the only model to consistently produce a FDR close to the nominal level, while CompDA was not able to avoid false discoveries if the effect was placed on the abundant cell type T4 (Figure E9). The superior performance of `cosmoDA` in this case was due to the fact the the data was simulated by an a-b power interaction model, which is not used by the other methods.

**Fig D1.**
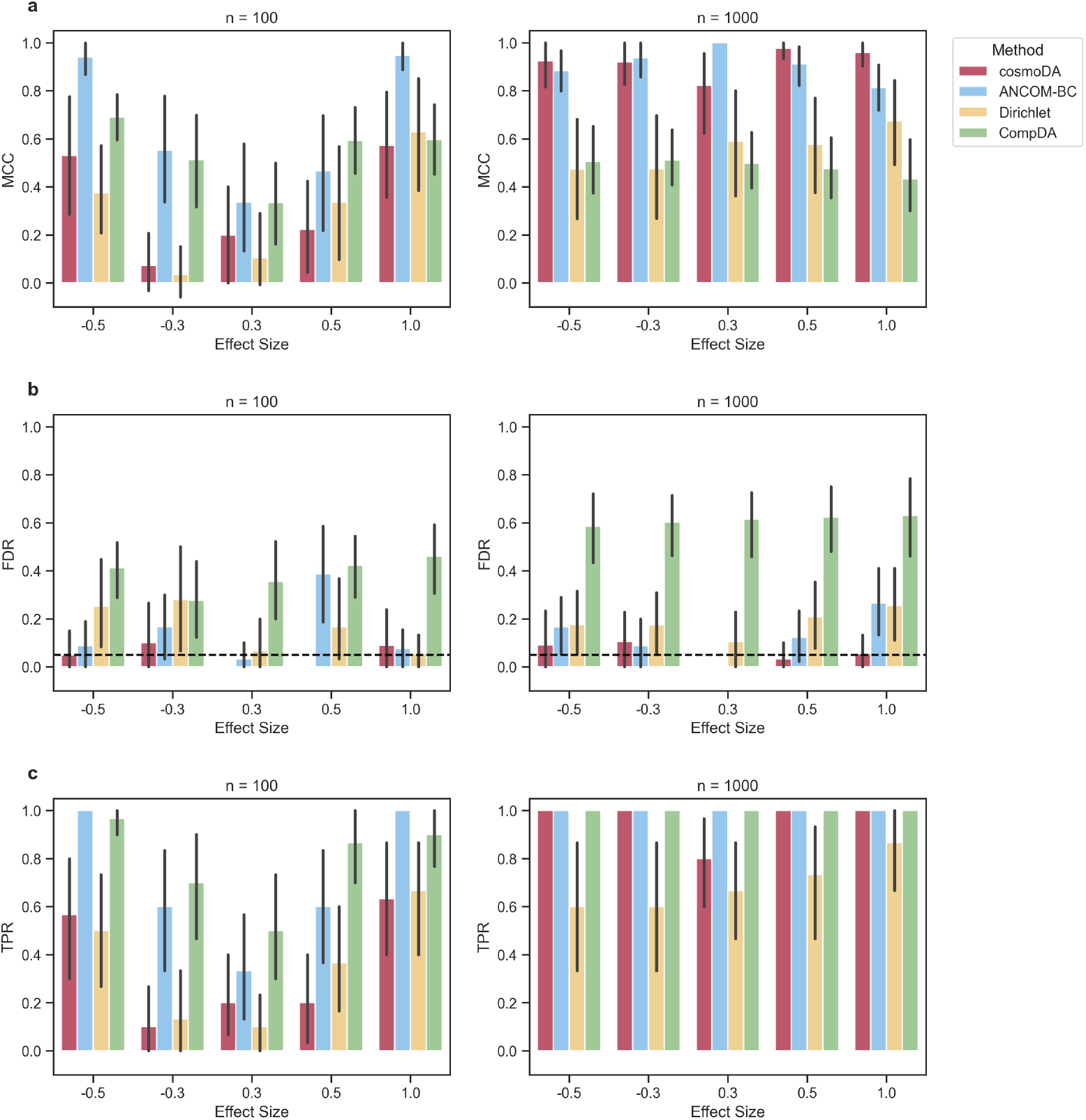
Performance comparison for recovering differentially abundant features across different scenarios, *K* = 0. (a) Matthews’ correlation coefficient. (b) False discovery rate. The dashed line shows the nominal FDR for all methods. (c) True positive rate (power).

## Appendix E Supplementary Figures

**Fig E2.**
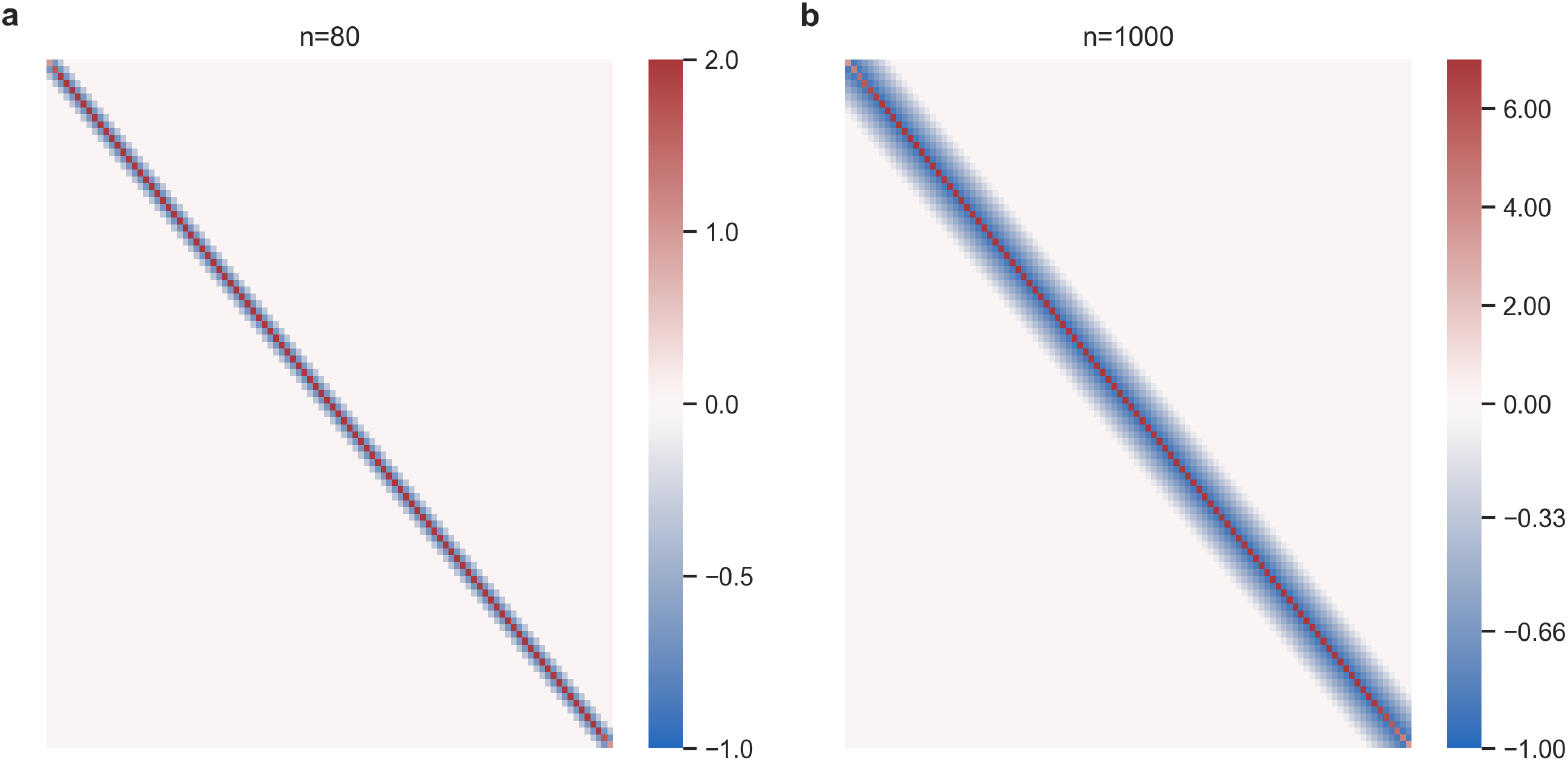
Interaction matrices used for data generation in the benchmark testing recovery of *K*(Section 3.1). (a) *n* = 80, (b) *n* = 1000.

**Fig E3.**
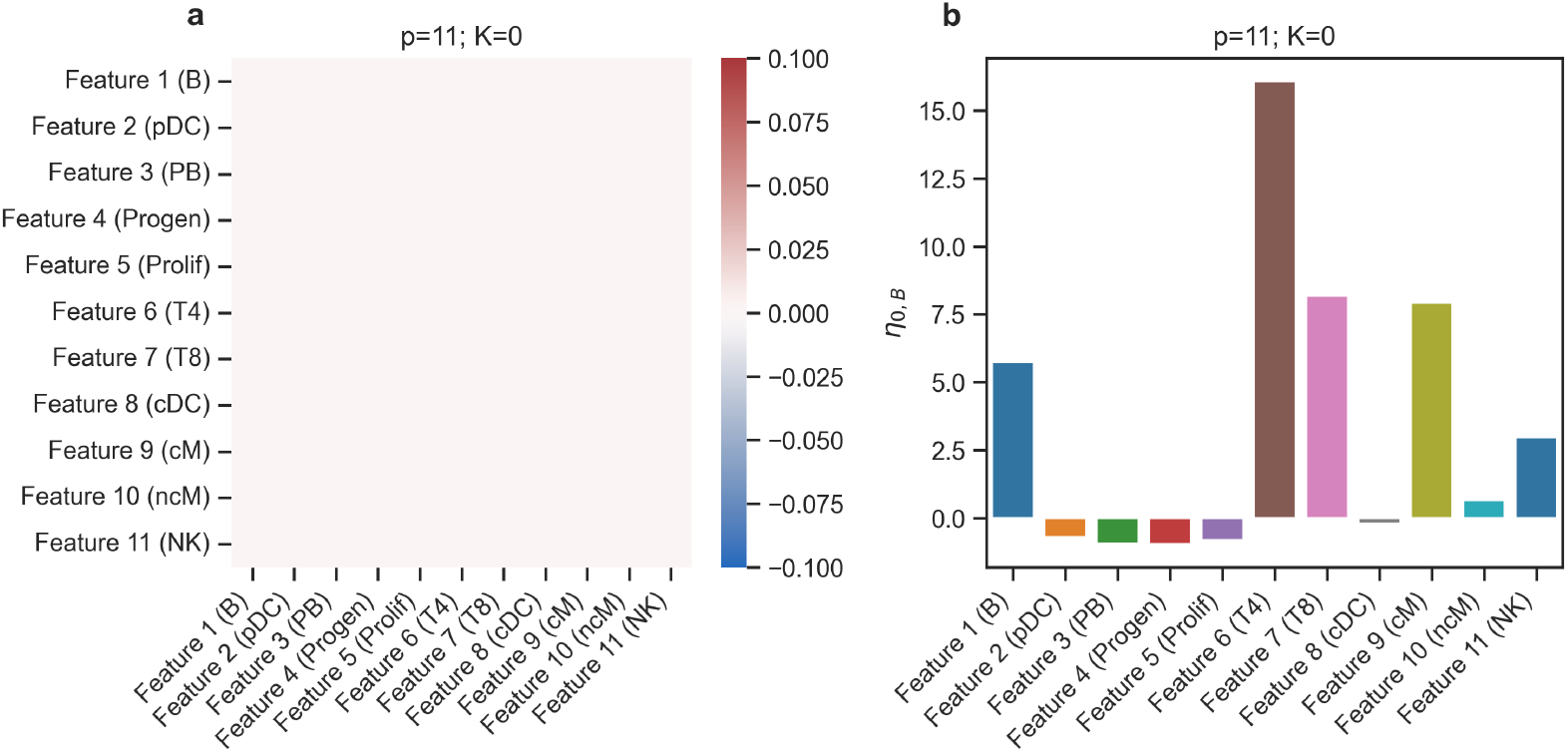
Data generation parameters used for the differential abundance testing benchmark (Section 3.2), *K* = 0. (a) Interaction matrix (***K***_*B*_). (b) Location vector (***η***_0,*B*_).

**Fig E4.**
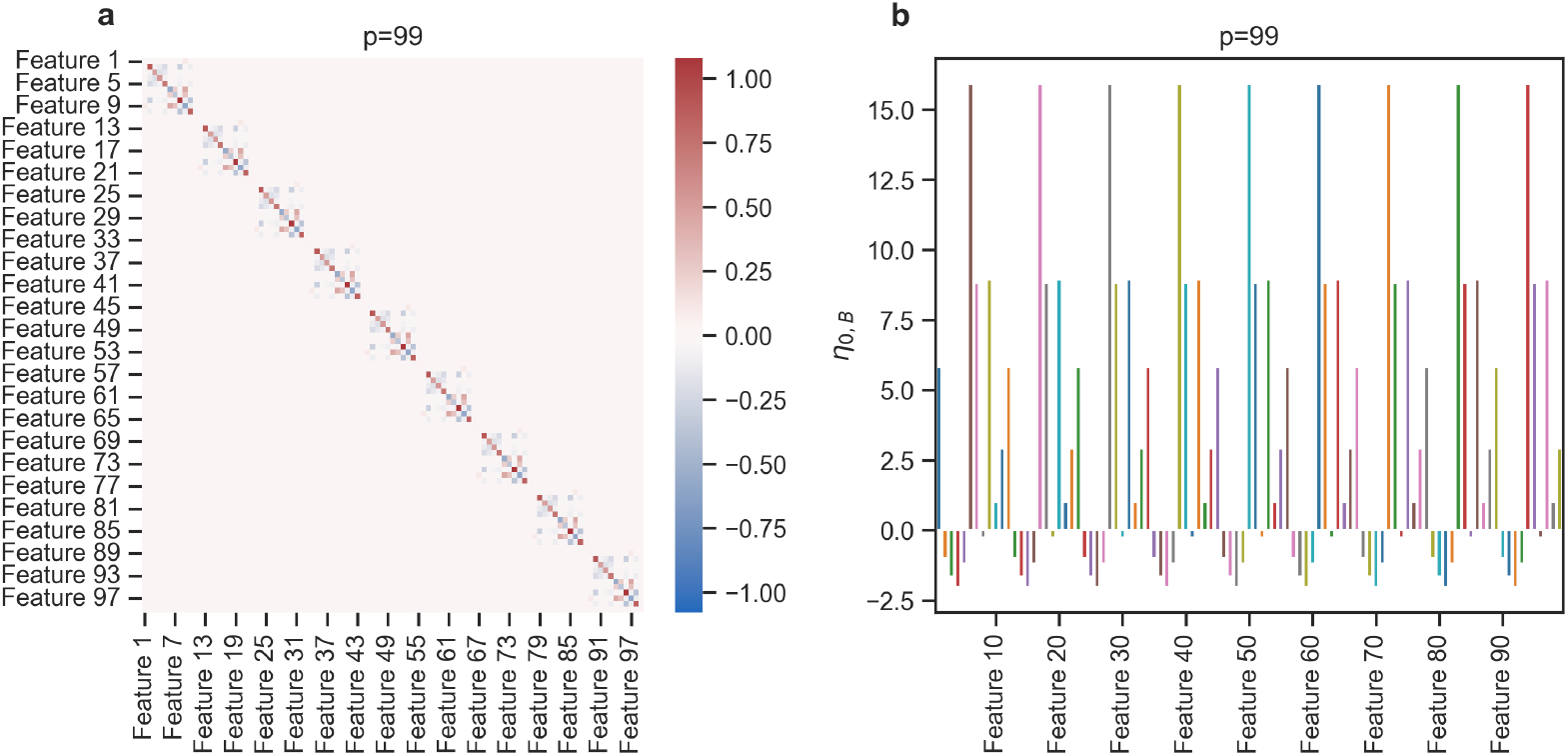
Data generation parameters used for the differential abundance testing benchmark (Section 3.2), *p* = 99. (a) Interaction matrix (***K***_*B*_). (b) Location vector (***η***_0,*B*_).

**Fig E5.**
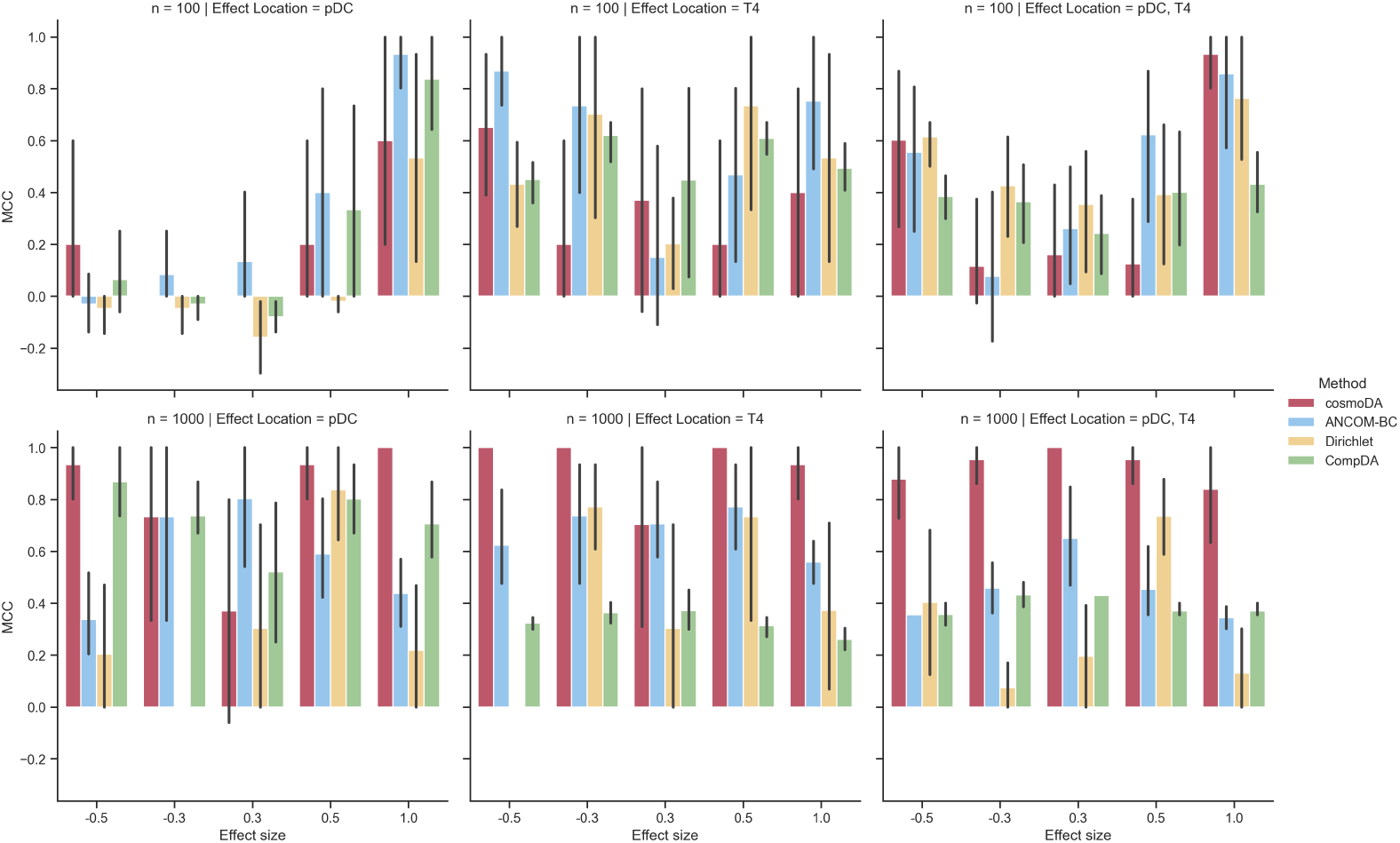
Detailed breakdown of Matthews’ correlation coefficient for the differential abundance testing benchmark (Section 3.2), *p* = 11.

**Fig E6.**
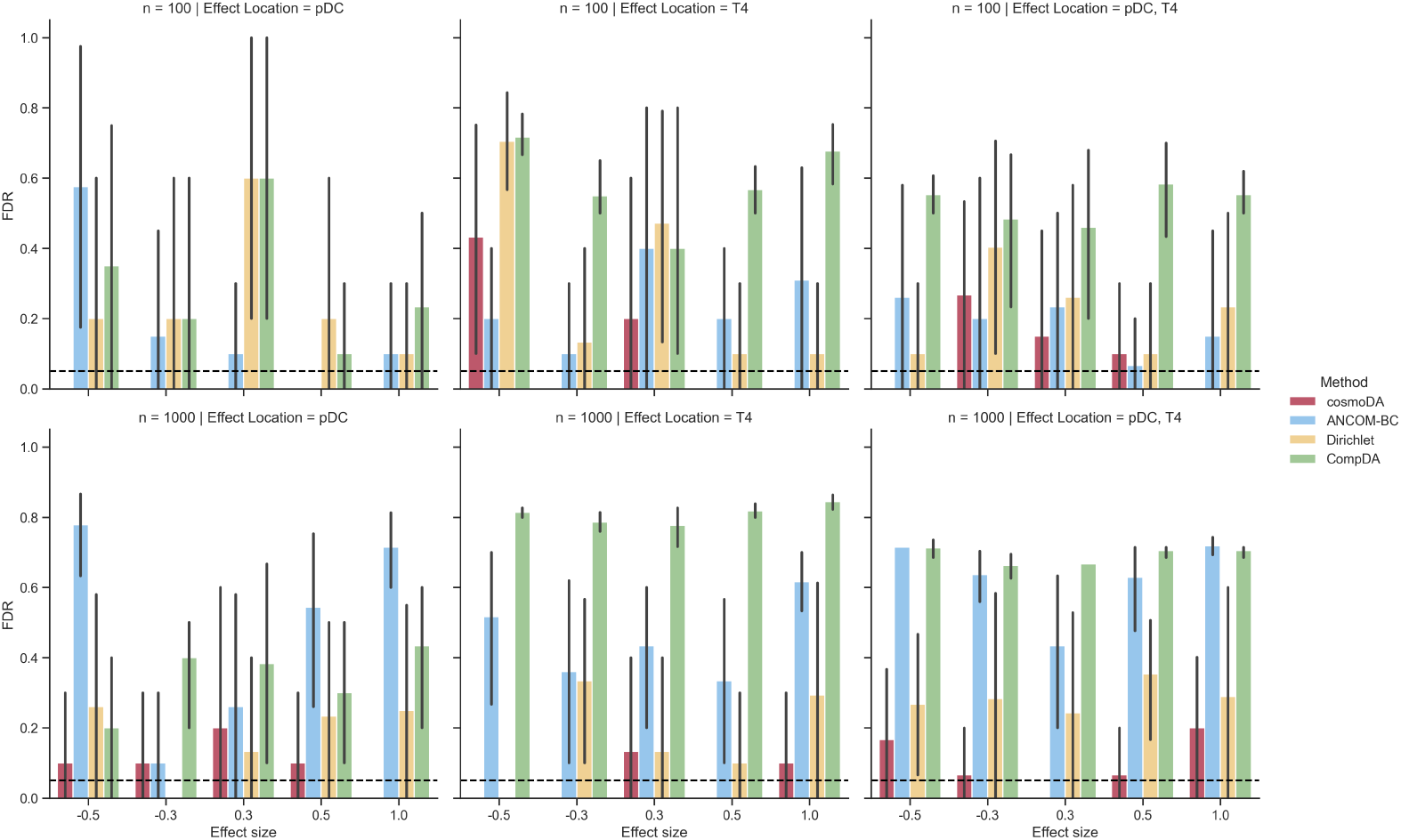
Detailed breakdown of false discovery rate for the differential abundance testing benchmark (Section 3.2), *p* = 11. The dashed lines denote the nominal FDR for all methods.

**Fig E7.**
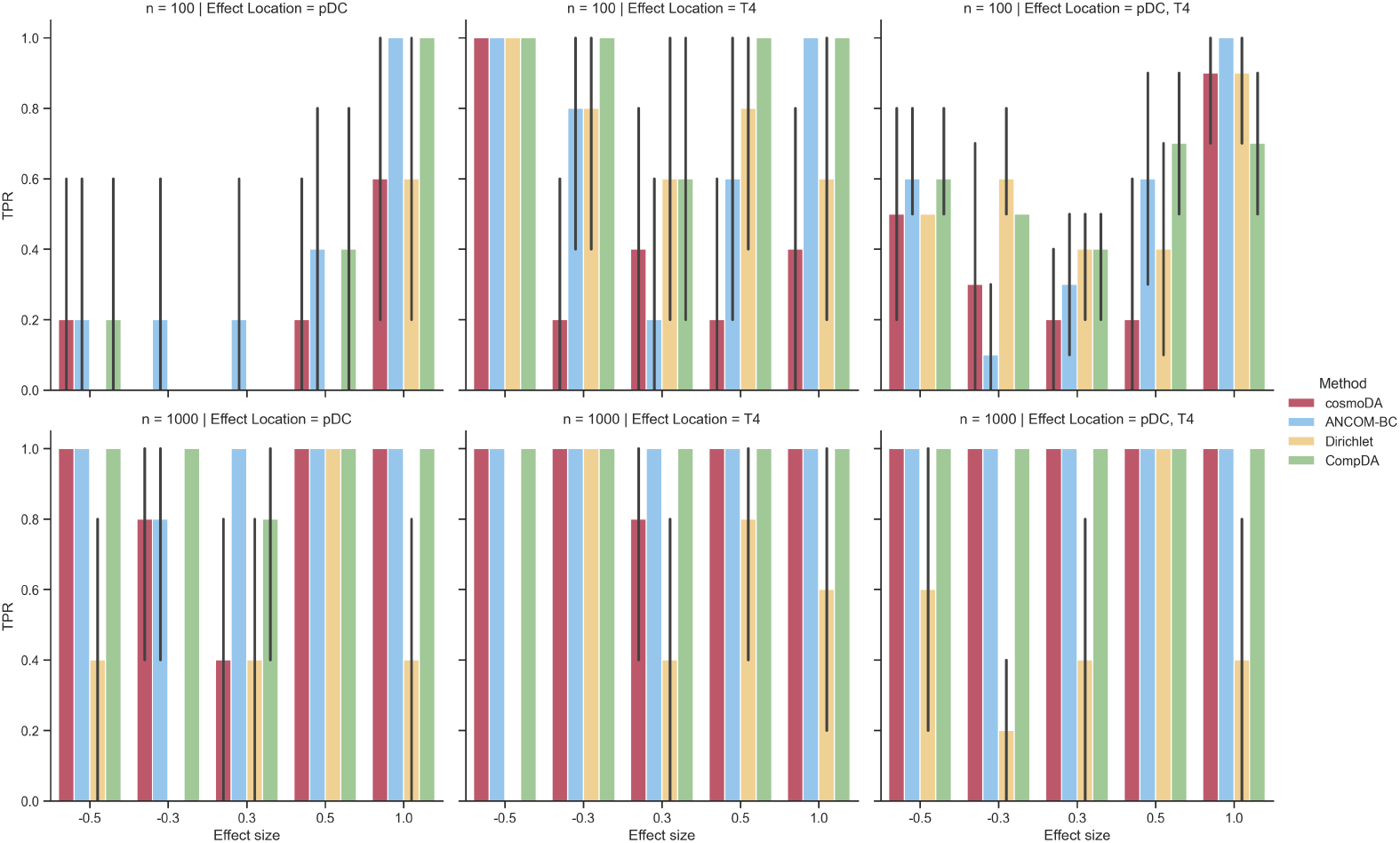
Detailed breakdown of power (true positive rate) for the differential abundance testing benchmark (Section 3.2), *p* = 11.

**Fig E8.**
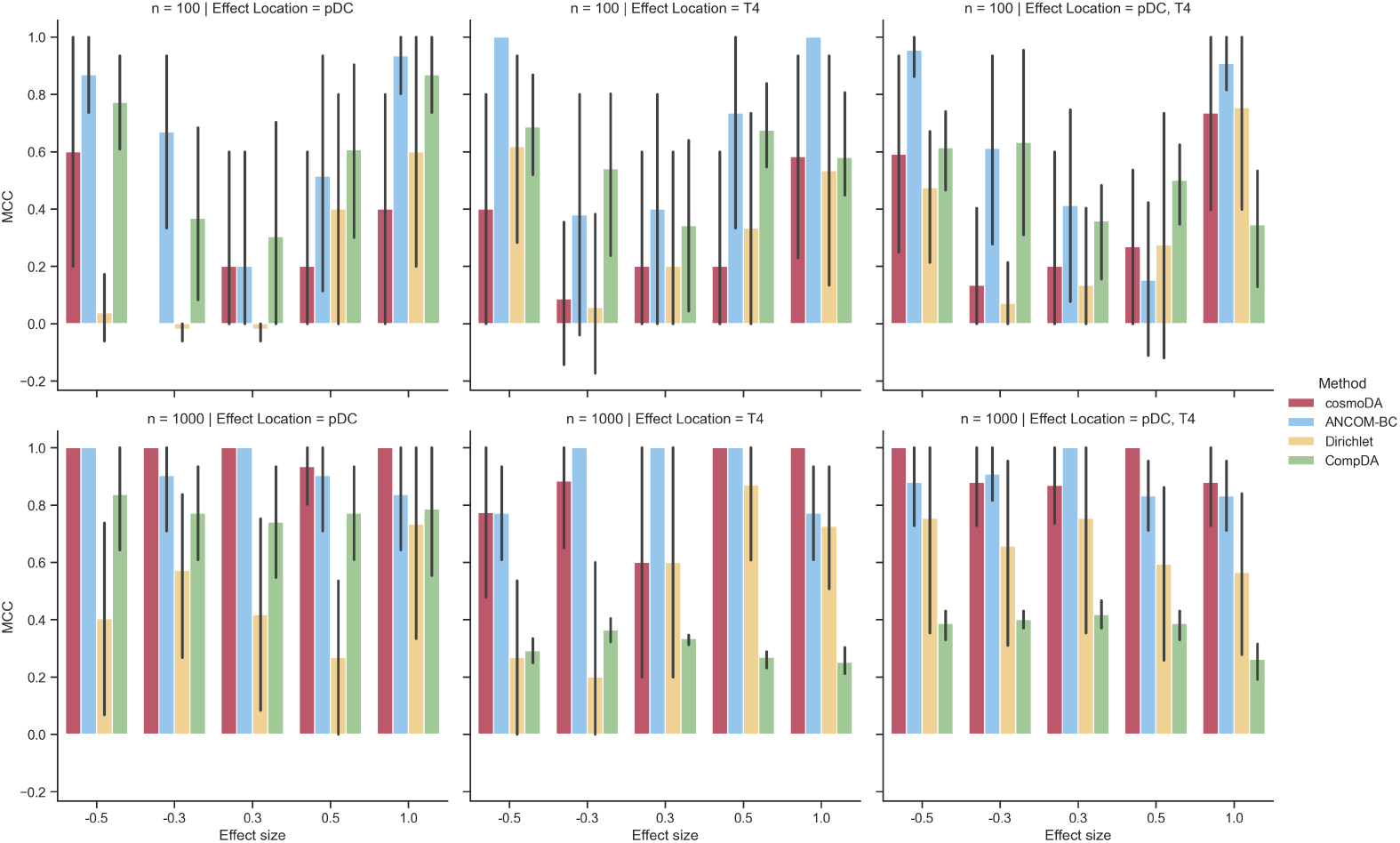
Detailed breakdown of Matthews’ correlation coefficient for the differential abundance testing benchmark (Section 3.2), *K* = 0.

**Fig E9.**
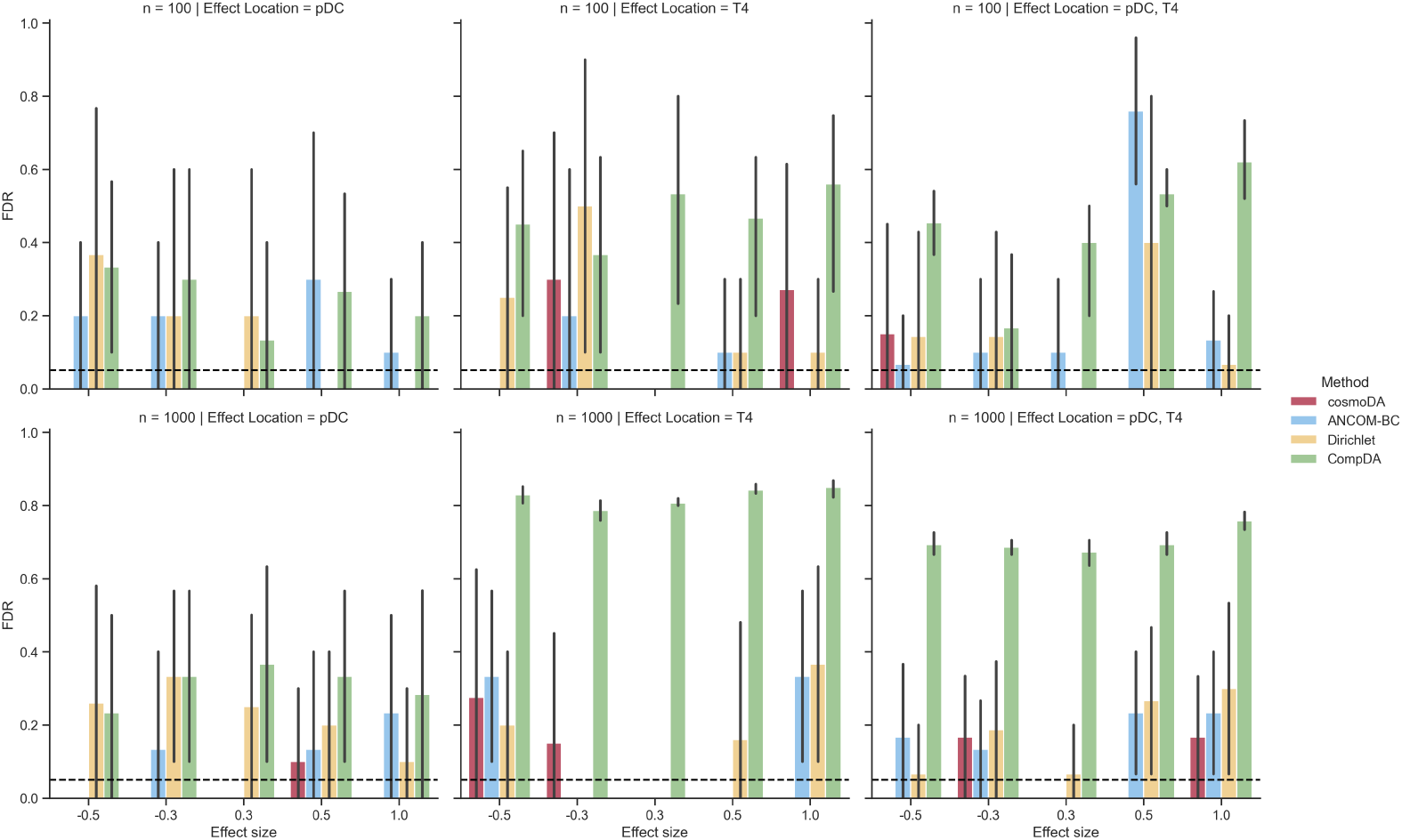
Detailed breakdown of false discovery rate for the differential abundance testing benchmark (Section 3.2), *K* = 0. The dashed lines denote the nominal FDR for all methods.

**Fig E10.**
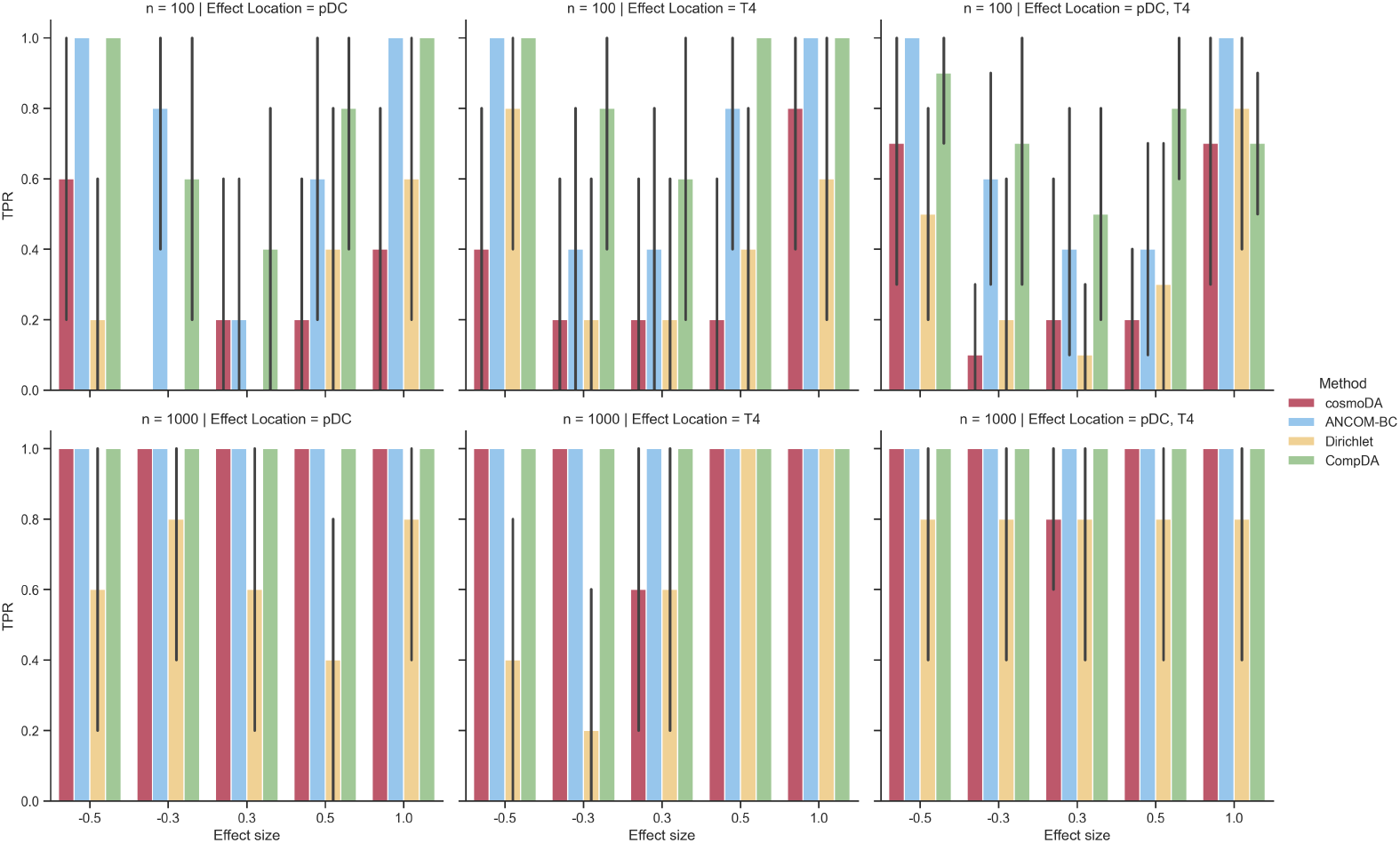
Detailed breakdown of power (true positive rate) for the differential abundance testing benchmark (Section 3.2), *K* = 0.

**Fig E11.**
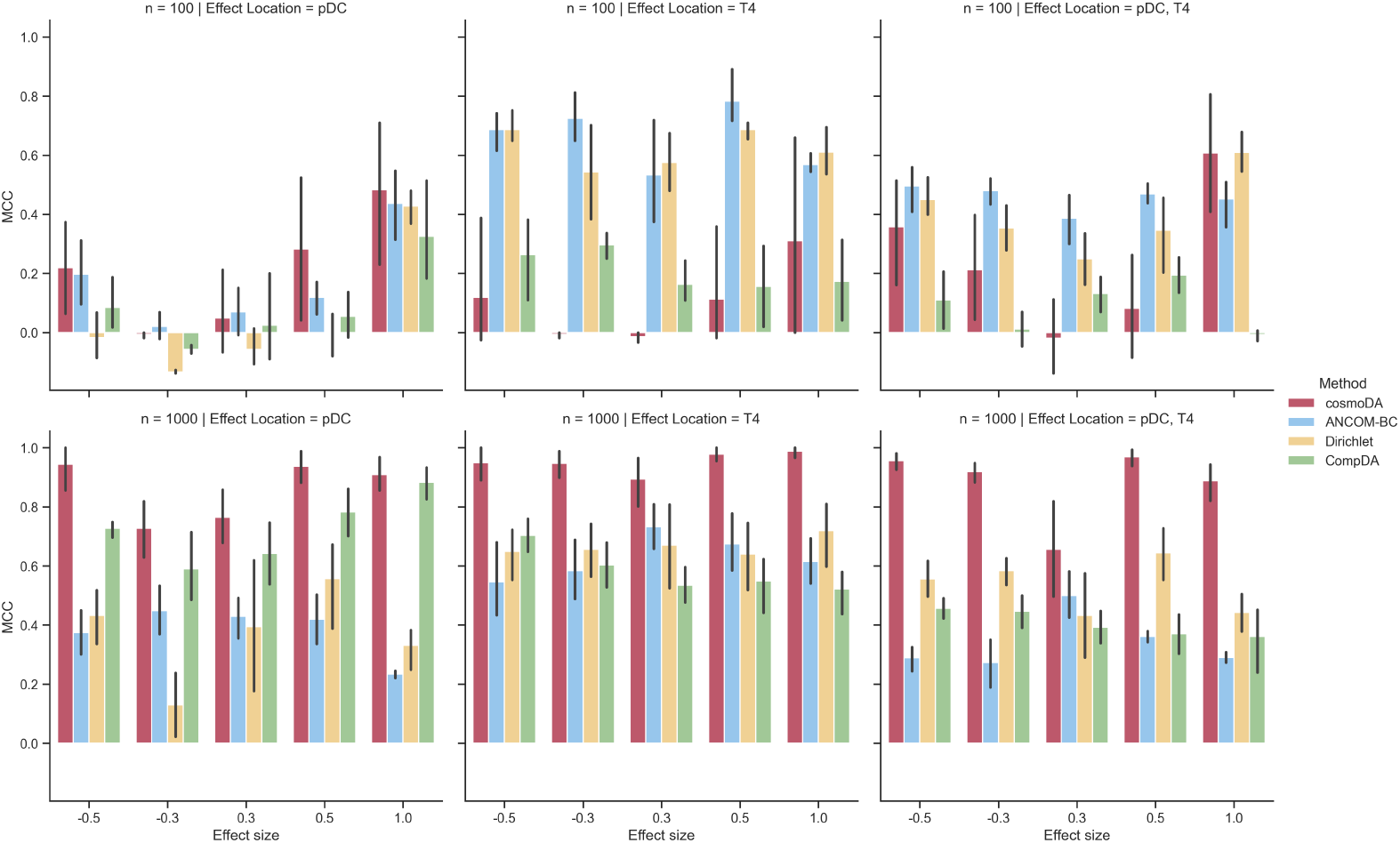
Detailed breakdown of Matthews’ correlation coefficient for the differential abundance testing benchmark (Section 3.2), *p* = 99.

**Fig E12.**
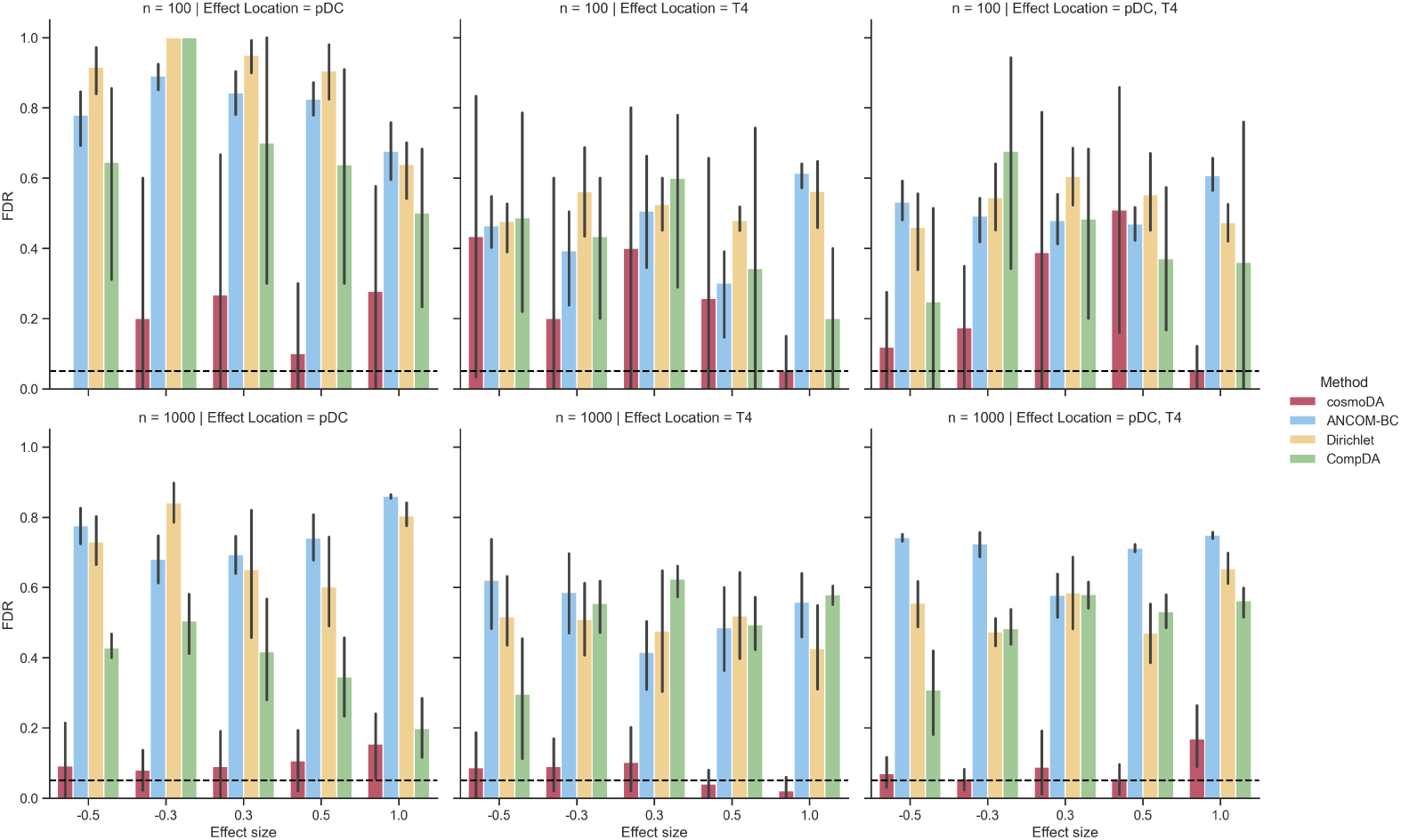
Detailed breakdown of false discovery rate for the differential abundance testing benchmark (Section 3.2), *p* = 99. The dashed lines denote the nominal FDR for all methods.

**Fig E13.**
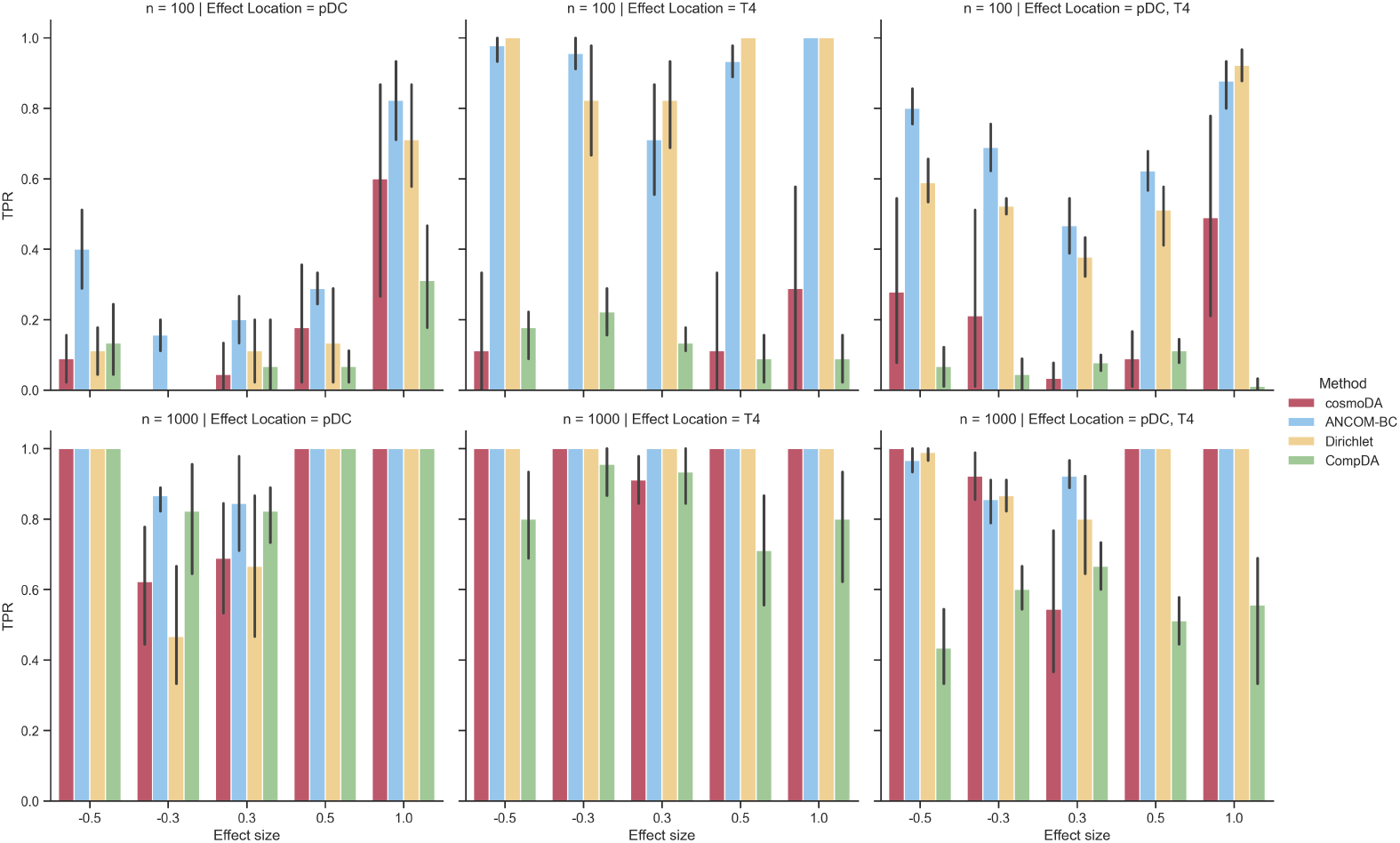
Detailed breakdown of power (true positive rate) for the differential abundance testing benchmark (Section 3.2), *p* = 99.

**Fig E14.**
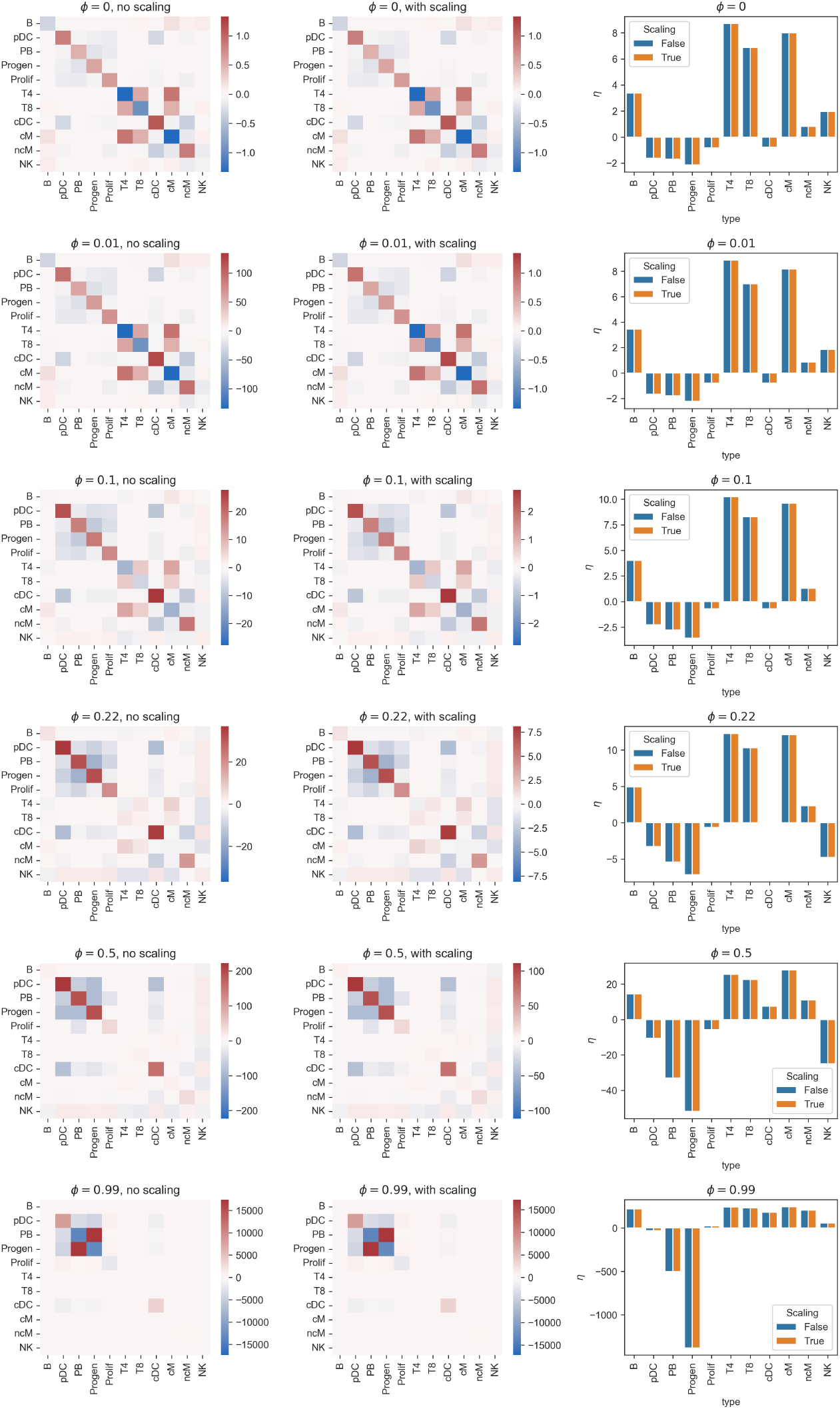
Impact of scaling Γ and *g* on the estimation of *K* and *η*. Results shown for the SLA scRNA-seq data Perez et al. (2022). Rows show selected values of the exponent *ϕ* in the power transformation. Left column: Values of ***K*** without scaling. Middle column: Values of ***K*** with scaling. Right column: Values of ***η*** with and without scaling.

**Fig E15.**
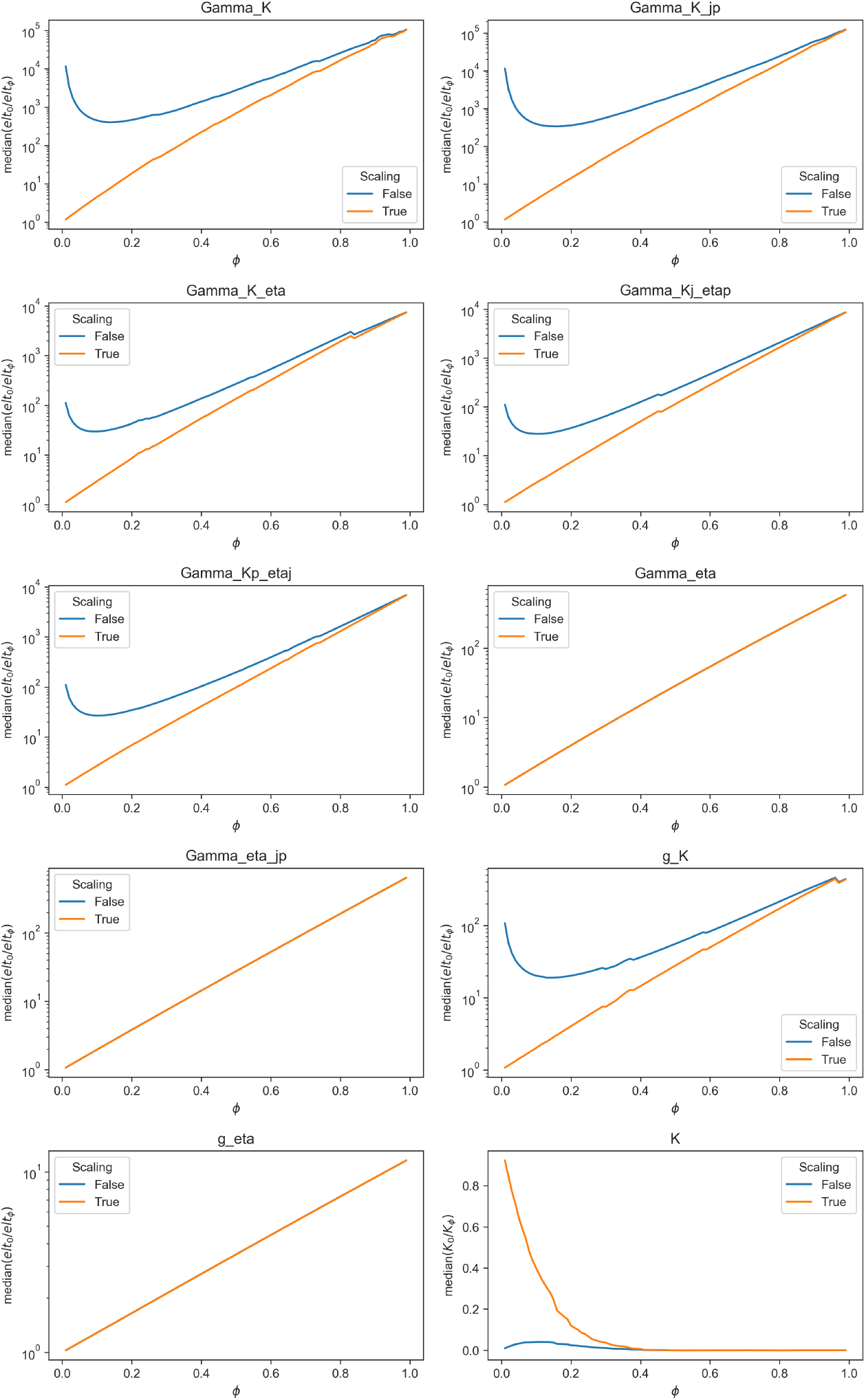
Impact of scaling factor for power transforms on the score matching parameters Γ and *g* and the interaction matrix *K*. All plots except bottom right show the median entry of *E*_*ϕ*=0_*/E*_*ϕ*=*ϕ*_^*′*^ for *E* being one of the score matching elements in Eq. 11. Bottom right: Same quantity for the estimated interaction matrix ***K***.

**Fig E16.**
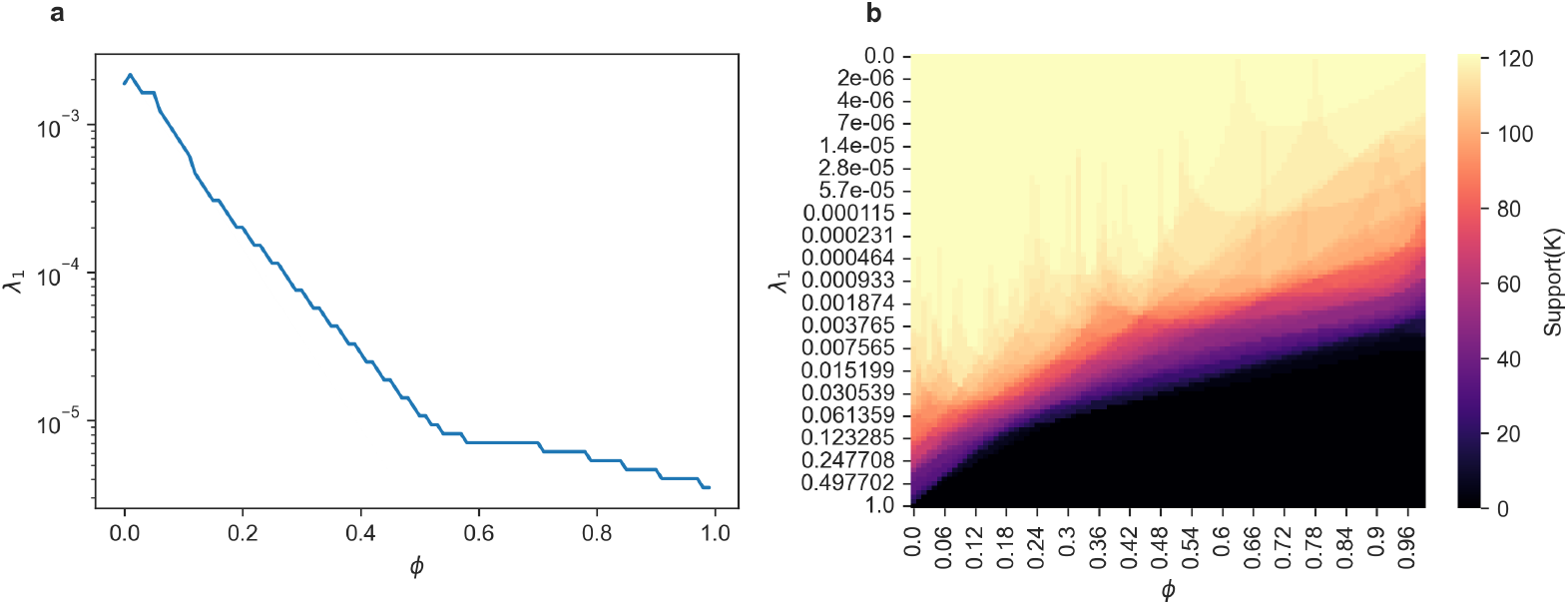
Relationship between power and regularization strength for the SLA scRNA-seq data Perez et al.(2022). (a) Value of *λ*_1_ selected through cross validation in relation to exponent *ϕ* of the power transform. (b) Number of nonzero entries in ***K*** for every *λ*_1_ and *ϕ*.

